# Chaotropic Ions Reshape the Cell Wall of the Obligate Halophile *Aspergillus atacamensis*: Insight from Solid-State NMR

**DOI:** 10.64898/2026.01.18.700186

**Authors:** Isha Gautam, Gisell Valdés Muñoz, Aswath Karai, Jean-Paul Latgé, Yordanis Pérez-Llano, Nina Gunde-Cimerman, Ramón Alberto Batista-García, Tuo Wang

**Author notes:** Correspondence and requests for materials should be addressed to T.W. and RAB-G.

## Abstract

Fungal survival in hypersaline environments requires exceptional adaptation of polysaccharide-based cell walls, yet the molecular principles underlying these adaptations remain largely unknown due to the extreme rarity of obligate halophilic fungi. *Aspergillus atacamensis* is an obligate halophile and chaotolerant fungus capable of growth at saturating NaCl concentrations and unusually high levels of MgCl_2_. Here, we used multidimensional solid-state NMR spectroscopy to investigate the molecular organization, hydration, and dynamics of cell wall polysaccharides in intact, uniformly ^13^C-labeled *A. atacamensis* cells grown under kosmotropic NaCl and chaotropic MgCl_2_ conditions. Under NaCl conditions, the rigid cell wall core was dominated by β-1,3-glucan and chitin across all salinities. Hyperosmotic NaCl induced thinner, dehydrated walls with increased polysaccharide mobility. In contrast, MgCl_2_ exposure resulted in marked remodeling of wall carbohydrates, including the emergence of chitosan, incorporation of mannan into the rigid phase, increased wall thickness, and enhanced hydration and dynamics. Together, these findings reveal fundamentally distinct polysaccharide remodeling strategies in response to kosmotropic versus chaotropic stress and establish a molecular framework for understanding fungal survival in extreme ionic environments.

**GRAPHICAL ABSTRACT:** 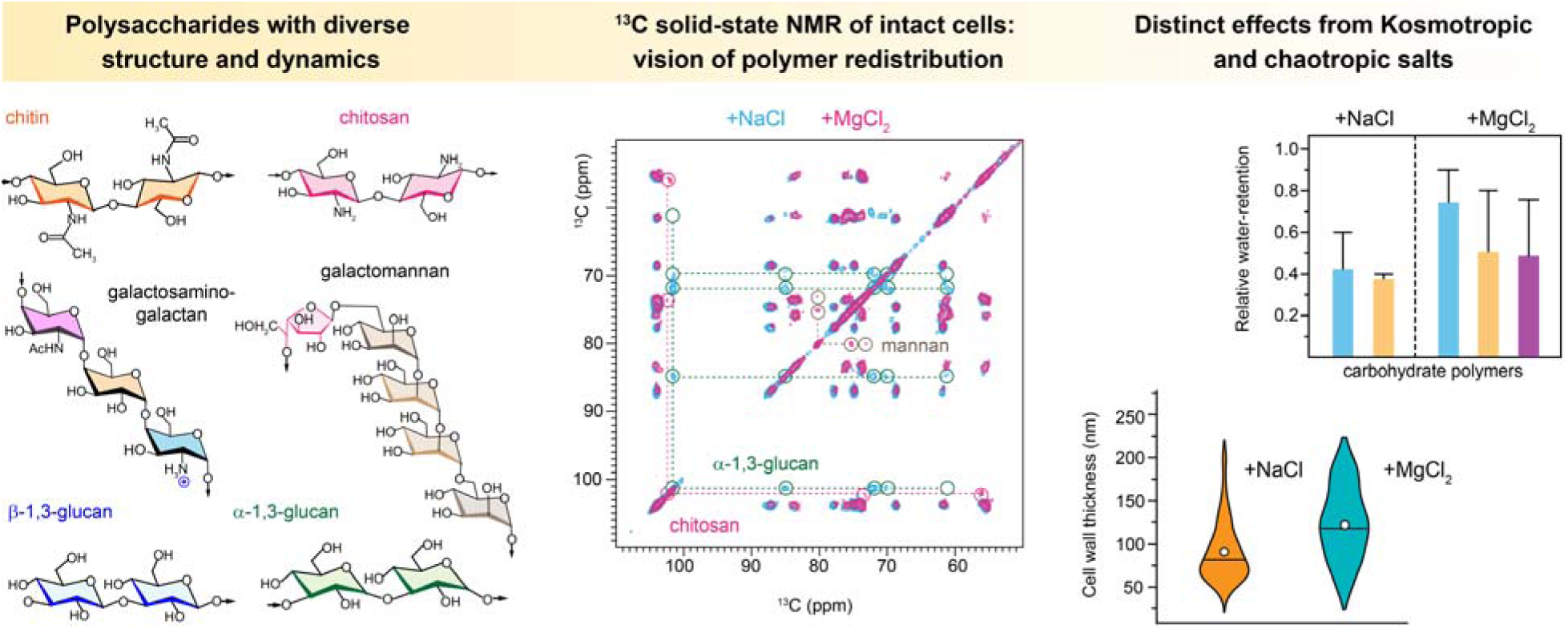

**HIGHLIGHTS:**

- Solid-state NMR reveals salt-dependent remodeling of fungal cell wall polysaccharides
- Kosmotropic and chaotropic salts drive distinct cell wall adaptation strategies
- NaCl stress promotes dehydration and compaction of the β-1,3-glucan-chitin framework
- MgCl_2_ induces chitosan emergence and alters mannan organization in the rigid phase
- Polysaccharide hydration and dynamics encode fungal adaptation to extreme environments

## INTRODUCTION

Among fungi, only a handful of species have been classified as obligate halophiles, including *Wallemia ichthyophaga*, *Wallemia muriae* (J. Zajc et al., 2014; Janja Zajc et al., 2013), *Aspergillus baarnensis* (= *Basipetospora halophila*), *Aspergillus salinarus* (Greiner et al., 2014), *Aspergillus destruens* (Sklenář et al., 2017), and, more recently, *Aspergillus salisburgensis* and *Aspergillus atacamensis* (Martinelli et al., 2017; Moreno-Perlin et al., 2023). The extreme rarity of obligate halophily in the fungal kingdom raises fundamental questions regarding the evolutionary adaptations that enable fungal survival in hypersaline environments (Gostinčar et al., 2022). Unlike halotolerant fungi, which can grow across a broad range of salinities, obligate halophiles require NaCl for growth and metabolic activity. This obligate dependence suggests specialized cellular mechanisms, particularly in the cell wall, which must maintain structural integrity while withstanding high ionic stress (Gunde-Cimerman et al., 2018; Moreno-Perlin et al., 2023). However, direct structural evidence of fungal cell wall remodeling in response to extreme salinity remains limited, mainly due to the lack of high-resolution techniques capable of examining cell wall architecture at atomic resolution without perturbing its native structure (Liyanage D Fernando et al., 2023; Kuncic et al., 2010).

*A. atacamensis* EXF-6660, initially isolated from a saltwater-exposed cave in the Coastal Range of the hyperarid Atacama Desert, represents a unique model for studying fungal halophily and chaotolerance (Martinelli et al., 2017; Moreno-Perlin et al., 2023). This species exhibits optimal growth between 1.5 M and 3.4 M NaCl and demonstrates remarkable tolerance to a variety of kosmotropic (stabilizing) and chaotropic (destabilizing) salts and solutes, including KCl, MgCl_2_, CaCl_2_, glycerol, and sorbitol, with limited tolerance to LiCl up to 1.7 M. Unlike many halotolerant fungi, *A. atacamensis* can reproduce at saturating NaCl concentrations, a feature observed in only a few extremophilic fungal species, such as *Aspergillus sydowii* and *W. ichthyophaga*, and in the highly halotolerant black yeast *Hortaea werneckii* (Gunde-Cimerman et al., 2018; Jiménez-Gómez et al., 2022; Moreno-Perlin et al., 2023; Janja Zajc et al., 2013). Furthermore, *A. atacamensis* exhibits metabolic versatility, including the ability to degrade polyaromatic compounds and benzene derivatives in the presence of 2.0 M NaCl, highlighting its potential for biotechnological applications (Moreno-Perlin et al., 2023).

One of the most intriguing aspects of *A. atacamensis* is its ability to thrive under chaotropic stress conditions, an area of fungal research that remains largely unexplored (González-Abradelo et al., 2025; Lever, 2016). In aqueous solutions, chaotropic agents such as MgCl_2_ disrupt macromolecular stability by interfering with water structure, while kosmotropic agents such as NaCl generally stabilize biomolecules (Cray et al., 2013; Hallsworth et al., 2007). However, at high concentrations, NaCl can also exert chaotropic effects, altering protein solubility, membrane integrity, and cellular architecture, similar to what has been observed in deep-sea brines and magnesium-rich hypersaline lakes (Hallsworth et al., 2007; Yakimov et al., 2015). While fungi such as *W. ichthyophaga*, *H. werneckii*, and *A. sydowii* demonstrate exceptional tolerance to a broad range of salts(J. Zajc et al., 2014), including high concentrations of MgCl_2_, the physiological, structural, and molecular responses of chaotolerant fungi to MgCl_2_ stress remain absolutely unknown(González-Abradelo et al., 2025). Notably, *A. atacamensis* has been documented to grow at MgCl_2_ concentrations (2.1 M) (Moreno-Perlin et al., 2023), far exceeding the previously established limit for microbial life (1.26 M) (Hallsworth et al., 2007), suggesting the presence of unique cellular adaptations that merit further investigation.

Understanding how extremophilic fungi restructure their cell walls in response to these extreme salinity conditions is a fundamental yet understudied aspect of microbial adaptation (Gunde-Cimerman et al., 2018). The fungal cell wall serves as the primary interface between the cell and its environment, playing a crucial role in maintaining osmotic balance, providing mechanical stability, and regulating stress responses (Ene et al., 2015; Garcia-Rubio et al., 2020). High salt concentrations are known to impact the biosynthesis and remodeling of key structural components, including glucans, chitin, and chitosan (Fuchs & Mylonakis, 2009), yet direct evidence of cell wall remodeling under extreme conditions remains scarce. Most studies on fungal osmoadaptation focus on compatible solutes and intracellular signaling pathways (Gostinčar et al., 2022; Gunde-Cimerman et al., 2018), with little emphasis on how the physical and biochemical properties of the cell wall are modified to withstand hypersaline stress.

Analyzing fungal cell wall polysaccharides presents significant challenges due to their complex composition and insolubility. Traditional biochemical methods involve cell wall isolation, solubilization, and chemical fractionation to characterize the structural components (Aimanianda et al., 2017; Bekirian et al., 2024; Black et al., 2023; Fontaine et al., 2000; Gravelat et al., 2013; J. P. Latgé, 2007). However, recent advances in high-resolution biophysical techniques, such as solid-state nuclear magnetic resonance (ssNMR), cryo-electron microscopy (CryoEM), and dynamic nuclear polarization (DNP), have revolutionized cell wall characterization by providing atomic-level insights without disrupting the native cellular architecture (I. Gautam et al., 2025; N. A. R. Gow & Lenardon, 2023; J.P. Latgé & Wang, 2022; Lenardon et al., 2020; Su et al., 2015). Multidimensional ssNMR spectroscopy, in particular, enables direct analysis of biomolecular interactions, hydration dynamics, and polymer organization in intact cells, making it a powerful tool for studying extremophilic fungi under stress conditions (Ghassemi et al., 2022; Polenova et al., 2015; Reif et al., 2021). Originally applied to plant biomass, bacterial cell wall and biofilms, and other extracellular matrices (Bougault et al., 2020; Byeon et al., 2025; Byeon et al., 2023; Rampratap et al., 2024; Reichhardt & Cegelski, 2014; Schanda et al., 2014; Takahashi et al., 2013; Vallet et al., 2024; Xue et al., 2024), these ssNMR methodologies have also been extended to fungal cell wall research, shedding light on the structural adaptations of fungal pathogens and industrially relevant species (Ankur et al., 2025; Chatterjee et al., 2015; Ehren et al., 2020; Jacob et al., 2025; Kang et al., 2018; Lamon et al., 2023; Lends et al., 2025; Safeer et al., 2023; Singh et al., 2025).

Despite these technological advancements, studies investigating how extremophilic fungi restructure their cell walls under hypersaline and chaotropic conditions are virtually nonexistent. To date, only a few reports have explored cell wall remodeling in obligate halophilic and chaotolerant fungi (Liyanage D Fernando et al., 2023). Given *A. atacamensis*’s ability to survive in conditions of extreme kosmotropic and chaotropic stress, this study provides the first molecular-level, high-resolution investigation of fungal cell wall architecture under extreme chaotropic conditions imposed by MgCl_2_.

In this study, ssNMR spectroscopy was utilized to analyze intact cells of *A. atacamensis*, detailing the changes in composition, dynamics, and hydration of cell wall polysaccharides under kosmotropic and chaotropic stress. Uniformly ^13^C, ^15^N-labeled *A. atacamensis* cells were subjected to hyperosmotic shock with NaCl and MgCl_2_ at both optimal and stress-inducing concentrations. While NaCl exposure had minimal impact on cell wall composition, MgCl_2_ induced significant structural adaptations, leading to the formation of thicker, more hydrated, and dynamic cell walls characterized by reduced chitin and mannan content and the emergence of chitosan. This study provides unprecedented detail on how an obligate halophilic and chaotolerant fungus remodels its cell wall in response to extreme environmental pressures. These findings offer fundamental insights into extremophilic fungal adaptations and inform the development of salt-tolerant fungal strains and biomaterials for biotechnological applications.

## MATERIALS AND METHODS

### Fungal strain, cultivation, and isotopic labeling

In this study, the extremophilic fungal strain *A. atacamensis* EXF-6660 was used as the model organism. This obligate halophile was isolated from NaCl brine flowing along the wall of a cave in the Atacama Desert, Chile (Martinelli et al., 2017). The strain was cultured and maintained on Potato Dextrose Agar (PDA) (Catalog # CM0139B, Thermo Fisher Scientific) supplemented with 1.7 M NaCl, which was determined to be the optimal salt concentration for its growth (Martinelli et al., 2017; Moreno-Perlin et al., 2023). Cultures were incubated at 28 °C for seven days to promote sporulation and mycelial development. Spores and mycelium were cryopreserved in liquid nitrogen and deposited in the Ex Microbial Culture Collection of the Infrastructural Centre Mycosmo at the Department of Biology, Biotechnical Faculty, University of Ljubljana, Slovenia.

For isotopic labeling, *A. atacamensis* was cultivated in 100 mL of liquid medium containing 20 g/L of uniformly ¹³C-labeled glucose (Catalog # CLM-1396-PK, Cambridge Isotope Laboratories) and 2 g/L of ¹ N-labeled ammonium nitrate (Catalog # NLM-712-PK, Cambridge Isotope Laboratories) as the sole carbon and nitrogen sources, respectively. To support fungal growth, the medium was supplemented with 10 g/L malt extract and 1 g/L peptone. A total of 200 mg of actively growing mycelium was aseptically transferred into sterilized liquid medium supplemented with the respective salts (as detailed in the next section) and incubated at 28°C in a shaking incubator (Corning, LSE) at 150 rpm.

### Saline stress treatments

To comprehensively evaluate the effects of salinity stress on the cell wall structure, fungal cultures were subjected to three distinct experimental conditions: (i) hyperosmotic NaCl stress, where cultures initially grown under 1.7 M NaCl, previously identified as the optimal salt concentration (Moreno-Perlin et al., 2023) for *A. atacamensis* EXF-6660, were transferred to 5.13 M saturated NaCl and maintained for 1 day and 4 days; (ii) optimal and suboptimal MgCl_2_ conditions, where cultures were grown under 1.5 M MgCl_2_ (optimal) and 0.5 M MgCl_2_ (suboptimal) (Moreno-Perlin et al., 2023) for 10 days; and (iii) conditions for comparative analysis, where cultures were maintained in their respective optimal conditions (1.7 M NaCl for 7 days or 1.5 M MgCl_2_ for 10 days) to enable direct comparisons between growth-optimized NaCl- and MgCl_2_-grown cultures. Additionally, optimal NaCl-grown cultures were extended to 10 days to enable direct comparison of the molecular level cell wall organization under long term growth in NaCl versus MgCl_2_ (1.7 M optimal concentration of NaCl versus 1.5 M optimal concentration of MgCl_2_, both grown for 10 days). It was not possible to implement cultures in the absence of salt because *A. atacamensis* is an obligate halophile and does not grow without salts, as our group has previously demonstrated (Martinelli et al., 2017; Moreno-Perlin et al., 2023). These conditions provided a framework for investigating fungal responses to varying osmotic stress levels and ion-specific growth environments, offering insights into the structural adaptations of the cell wall under different salinity regimes.

### Transmission electron microscopy (TEM) imaging

We analyzed cell wall thickness in *A. atacamensis* EXF-6660 using transmission electron microscopy (TEM) under different osmotic conditions. These included cultures grown under optimal NaCl conditions (1.7 M, 7 days) and optimal MgCl_2_ conditions (1.5 M, 10 days), as well as those exposed to hyperosmotic NaCl stress (5.13 M) for 1 day and 4 days. Additionally, we examined fungal samples grown under suboptimal MgCl_2_ conditions (0.5 M, 10 days) to assess the structural adaptations of the cell wall under varying salinity levels.

Fungal mycelia were fixed in a solution containing 2.5% glutaraldehyde, 2% paraformaldehyde, and 0.1 M cacodylate buffer. To maintain cellular integrity and prevent shrinkage, the samples were embedded in 2% agarose. A secondary fixation step was performed using 0.1 M osmium tetroxide. Dehydration was performed using a graded acetone series, followed by infiltration with epoxy resin-acetone mixtures at 25:75, 50:50, and 75:25 ratios. Samples were left overnight in the 75:25 resin-acetone mixture, then transferred to 100% resin for two days, with multiple resin exchanges. Final embedding was achieved by curing the resin blocks at 70°C. Ultrathin sections were prepared using a LEICA EM UC7 microtome, stained with 1% uranyl acetate and lead acetate, and mounted on carbon-coated grids. TEM imaging focused on hyphal cross-sections, with particular emphasis on cell wall structure.

Cell wall thickness was measured for each sample group, with 105-120 measurements per condition (**Table S1**). Image analysis was performed using ImageJ software (version 1.54g). Differences in cell wall thickness across the studied conditions were statistically analyzed using an unpaired two-tailed Student’s t-test with a significance threshold of *p* < 0.05. Statistical calculations were conducted in Microsoft Excel 365, and color-coded violin plots were generated using OriginPro 2021 to visualize the distribution of cell wall thickness measurements under different experimental conditions.

### 13C solid-state NMR

High-resolution ssNMR experiments were performed using a Bruker Avance Neo 800 MHz NMR spectrometer located at the Max T. Roger NMR Facility at Michigan State University (MI, USA). For ^13^C detection, a 3.2 mm HCN probe was employed with a magic-angle spinning (MAS) frequency of 15 kHz. Approximately 35-40 mg of sample was packed into the rotor. The experimental temperatures were maintained between 277 K and 298 K. The ^13^C chemical shifts were externally referenced by calibrating the CH_2_ peak of adamantane to 38.48 ppm, which was then used to establish the spectral reference for the spectra collected on fungal samples. The radiofrequency pulse field strengths applied during the experiments were 71.4-83.3 kHz for ^1^H hard pulses, decoupling, and cross-polarization, and 50-62.5 kHz for ^13^C pulses. Datasets on dynamics and hydration were conducted using a Bruker Avance Neo 400 MHz spectrometer equipped with a 3.2 mm triple-resonance HCN probe. Water-edited experiments were carried out at 277 K, while dynamics were measured at 298 K. Further details regarding the experimental conditions are provided in **Table S2**.

### Resonance assignment of carbohydrate signals

Multiple 2D ssNMR experiments were employed to investigate the molecular structure of the *A. atacamensis* cell wall. The through-bond ^13^C connectivity of mobile molecules was tracked using the refocused J-INADEQUATE experiment (Cadars et al., 2007; Lesage et al., 1999), utilizing a direct polarization (DP) scheme and 2-second recycle delays (**Fig. S1**). The J-evolution period consisted of four delays of 2.3 ms each, optimized to enhance the intensity of carbohydrate signals. Rigid molecules (**Fig. S2**) were detected using a CORD sequence that relies on cross-polarization (CP) for initial polarization creation, with a mixing time of 53 ms at 15 kHz MAS (Hou et al., 2013; Lu et al., 2015). Analysis of these 2D data allowed resonance assignments of carbon sites in carbohydrates, which were cross-referenced with chemical shift data from the Complex Carbohydrate Magnetic Resonance Database and recent studies (Kang et al., 2020; Zhao et al., 2023). The complete assignments are provided in **Table S3**.

### Estimation of carbohydrate composition

The molar composition of carbohydrates in each sample was estimated by analyzing peak volumes in 2D ^13^C-^13^C spectra obtained using 53 ms CP-CORD and DP refocused J-INADEQUATE experiments, which represented rigid and mobile polysaccharides, respectively (**Table S4, 5**). Peak volumes were extracted using the integration function of Bruker TopSpin software (version 4.1.4). To minimize uncertainty caused by spectral crowding, only well-resolved signals were considered for compositional analysis. Quantification in CORD spectra involved averaging the resolved cross-peaks for each carbohydrate, whereas in INADEQUATE spectra, only clearly defined spin connections were used. The NMR peaks used for compositional analysis, along with their resonance assignments and corresponding peak volumes, are provided in **Table S4, 5**. Relative abundance was determined by normalizing the sum of integrals to their respective counts. Standard errors were calculated based on the standard deviation of the integrated peak volume divided by the total number of cross-peaks. The overall standard error was derived from the square root of the sum of the squared standard errors for each polysaccharide, as described in recent studies (Cheng et al., 2024; Isha Gautam et al., 2025).

### Dynamics and hydration profile of carbohydrates

All experiments investigating polymer hydration and dynamics were conducted on a Bruker Avance Neo 400 MHz NMR spectrometer at 277 K and 298 K for hydration and dynamics experiments, respectively. To assess water accessibility of polysaccharides, 1D ^13^C and 2D water-edited ^13^C-^13^C correlation spectra were measured (Ader et al., 2009; Luo & Hong, 2010; White et al., 2014). A hard 90° pulse on the ^1^H channel was applied to excite all protons, followed by a ^1^H-T_2_ relaxation filter to suppress magnetization of proton resonances with shorter ^1^H-T_2_ values. The ^1^H-T_2_ relaxation filter was set to 1.3 ms × 2 for most samples, except for the hyperosmotic stress condition (1-day sample, 1.2 ms × 2), reducing polysaccharide intensity to less than 5% while preserving approximately 80% of water magnetization. Water ^1^H polarization was then transferred to nearby biomolecules during a ^1^H-^1^H mixing period, followed by 1-ms CP for high-resolution ^13^C detection. The ^1^H mixing time varied from 0 to 100 ms for a series of 1D water-edited spectra, and it was set to 4 ms for the 2D water-edited experiment with 50 ms of DARR mixing. Build-up curves for 1D spectra were obtained by plotting relative intensities as a function of the square root of the ^1^H mixing time (**Fig. S3**). For 2D spectra, intensity ratios (S/S_0_) between the water-edited spectrum (S) and the non-edited control spectrum (S_0_) were determined for each resolved carbon site, reflecting the degree of water retention (**Table S6**).

The dynamics of cell wall polysaccharides were probed using ^13^C spin-lattice (T_1_) relaxation times, determined through 1D Torchia CP correlation spectra with variable z-filter periods ranging from 0.1 μs to 12 s (**Fig. S4 and Table S7**). Additionally, ^1^H-T_1_ relaxation times were measured using the Lee-Goldburg (LG) spin-lock sequence (Duan & Hong, 2024; Huster et al., 2001; Van Rossum et al., 2000) with ^1^H spin-lock times ranging from 0.1 µs to 19 ms (**Fig. S5 and Table S7**). This sequence suppressed ^1^H-^1^H dipolar couplings during the spinlock and CP periods, providing carbohydrate-specific insights into polymer dynamics. Peak intensities were normalized to the number of scans and fit using a single-exponential decay equation to obtain the relaxation time constants for different carbon sites.

## RESULTS AND DISCUSSION

### Restructured rigid core of *A. atacamensis* cell walls under NaCl stress

We used *A. atacamensis* was used as a model obligate halophile to investigate mycelial characteristics under optimal sodium chloride conditions of 1.7 M NaCl. Cultures were grown for 7 days, corresponding to a time point within the exponential growth phase that yielded sufficient biomass for ssNMR analysis (Moreno-Perlin et al., 2023). Hyperosmotic shock conditions of 5.13 M NaCl were also examined, with samples collected after 1 day to represent the lag phase and after 4 days to represent the exponential phase. In addition, to enable direct comparison with data obtained under optimal MgCl_2_ conditions, which supported slower growth and required a 10-day culture to obtain sufficient material for ssNMR analysis, a corresponding sample grown in 1.7 M NaCl for 10 days was also prepared.

Solid-state NMR analysis of the four *A. atacamensis* samples revealed that β-1,3-glucan and chitin (**Fig. 1A**) dominate the rigid structural domain of the fungal cell wall under all tested growth conditions, as evidenced by their prominent signals in 1D ^13^C CP spectra (**Fig. 1B**) and 2D ^13^C-^13^C correlation spectra (**Fig. 1C**). Quantitative peak volume analysis demonstrated that the ratio of β-1,3-glucan to chitin remained consistent at approximately 40:60 in samples cultured for 1 to 4 days under hyperosmotic conditions, where a stable cell wall architecture may be required regardless of growth phase, but varied substantially in cultures grown in 1.7 M NaCl for 7 to 10 days, where cell wall remodeling occurs under optimal growth conditions (**Fig. 1D**).

**Figure 1.**
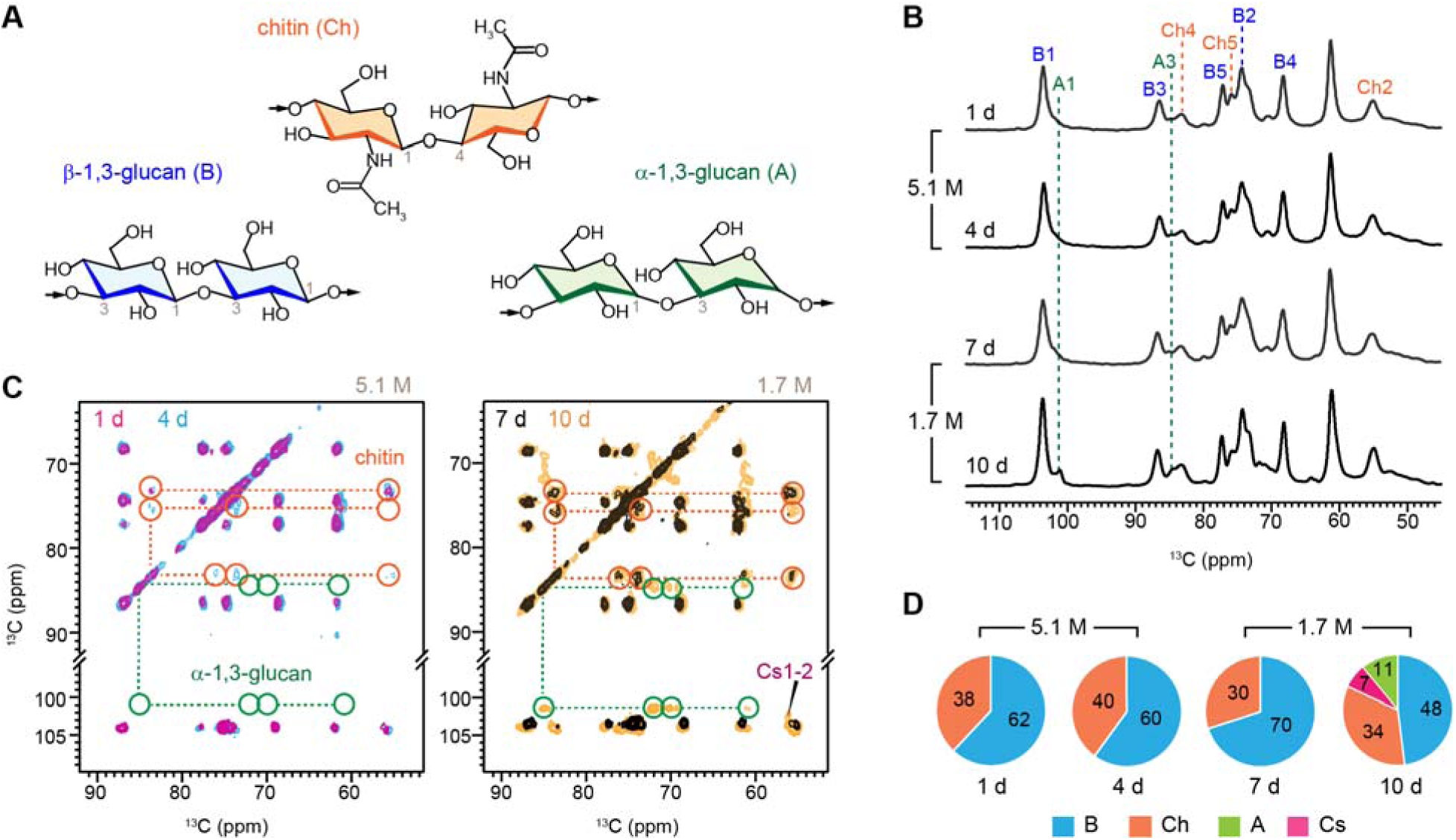
Carbohydrate composition in *A. atacamensis* under varied NaCl concentrations. (**B**) Simplified structural representation of glucans and chitin. (**C**) 1D ^13^C CP spectra of *A. atacamensis* cultured under different conditions. The 1-day and 4-day cultures were grown in 5.13 M NaCl under hyperosmotic conditions, while the 7-day and 10-day cultures were grown in 1.7 M NaCl, which is considered optimal for growth. (**D**) Overlay of 2D ^13^C-^13^C 53-ms CORD spectra collected on *A. atacamensis* grown under hyperosmotic conditions for 1 day (magenta) and 4 days (cyan), highlighting chitin development in the 4-day sample and the absence of α-1,3-glucan in both samples. On the right, the overlay of spectra collected from 7-day (black) and 10-day (yellow) samples highlights the emergence of α-1,3-glucans during extended growth under optimal condition. (**E**) Pie charts showing the molar composition of rigid carbohydrates based on intensity analysis of 2D ^13^C CORD spectra. B: β-1,3-glucan; Ch: chitin; A: α-1,3-glucan; Cs: chitosan.

Interestingly, the rigid domain of the cell wall also displayed α-1,3-glucan in the sample grown under optimal conditions (1.7 M NaCl) for 10 days (**Fig. 1B, C**). In this extended growth condition, the presence of chitosan was confirmed by the cross-peak between carbon 1 and carbon 2 (Cs1-2), indicating partial deacetylation of chitin at this stage (**Fig. 1C**). While the 7-day-old sample had a relatively simple rigid core, with a β-1,3-glucan-to-chitin ratio of 70:30, the 10-day-old cell wall displayed increased complexity (**Fig. 1D**). This sample comprised 48% β-1,3-glucan, 34% chitin, 11% α-1,3-glucan, and 7% chitosan, reflecting significant structural remodeling during prolonged growth under optimal conditions.

To complement the NMR structural data, we conducted transcriptomic analyses to examine the transcriptional response of *A. atacamensis* to NaCl-induced hyperosmotic stress and to relate cell wall remodeling to underlying molecular pathways (**Supplementary Methods**). Glucan-related genes displayed dynamic regulation during 1-4 days of high-salinity exposure compared to optimal 7-day growth (**Fig. S6**), with upregulation of β-1,3-glucan biosynthetic genes such as β-1,3-glucanosyltransferase and β-1,3-glucan synthase (Hu et al., 2023). In contrast, chitin metabolism genes and mannose-related genes were downregulated, a pattern also reported for *A. sydowii* under extreme salinity (Jiménez-Gómez et al., 2022). Although substantial transcriptional regulation of these cell wall biosynthesis genes was observed, these changes did not fully align with structural observations, echoing similar discrepancies reported in *A. sydowii*, where transcriptomic variation did not directly correspond to cell wall composition changes (Liyanage D Fernando et al., 2023). This supports the hypothesis that halophilic fungi maintain a stabilized cell wall architecture through buffering mechanisms that operate independently of transcription, potentially involving post-transcriptional or post-synthetic controls of polymer assembly.

### Incorporation of chitosan and mannan into the rigid cell wall under MgCl_2_ cultivation

The responses of *A. atacamensis* to magnesium chloride (MgCl_2_) differ significantly from those observed under sodium chloride (NaCl). Chitosan signals were consistently detected in 10-day-old samples cultured under both stressed (0.5 M) and optimal (1.5 M) MgCl_2_ concentrations. These signals, corresponding to chitosan carbon 1 and carbon 2 (Cs1 and Cs2), were evident in 1D ^13^C spectra (**Fig. 2A**) and in cross-sections of 2D ^13^C-^13^C CORD correlation spectra (**Fig. 2B**). Additionally, signals associated with α-1,2-linked mannopyranose units were observed in the 2D ^13^C-^13^C CORD spectra of both samples (**Fig. 2C**). These mannopyranose residues, the primary components of the galactomannan (GM) backbone in *Aspergillus* species, are typically found in the mobile fraction of the cell wall (Chakraborty et al., 2021; J.P. Latgé & Wang, 2022). Their presence in the CORD spectra that preferentially detect rigid molecules, indicates that portions of the galactomannan backbone have interacted with chitin microfibrils, thereby rigidifying the polymer backbone. This provides clear evidence of the differences in cell wall organization between mycelia grown under NaCl and MgCl_2_ exposure.

**Figure 2.**
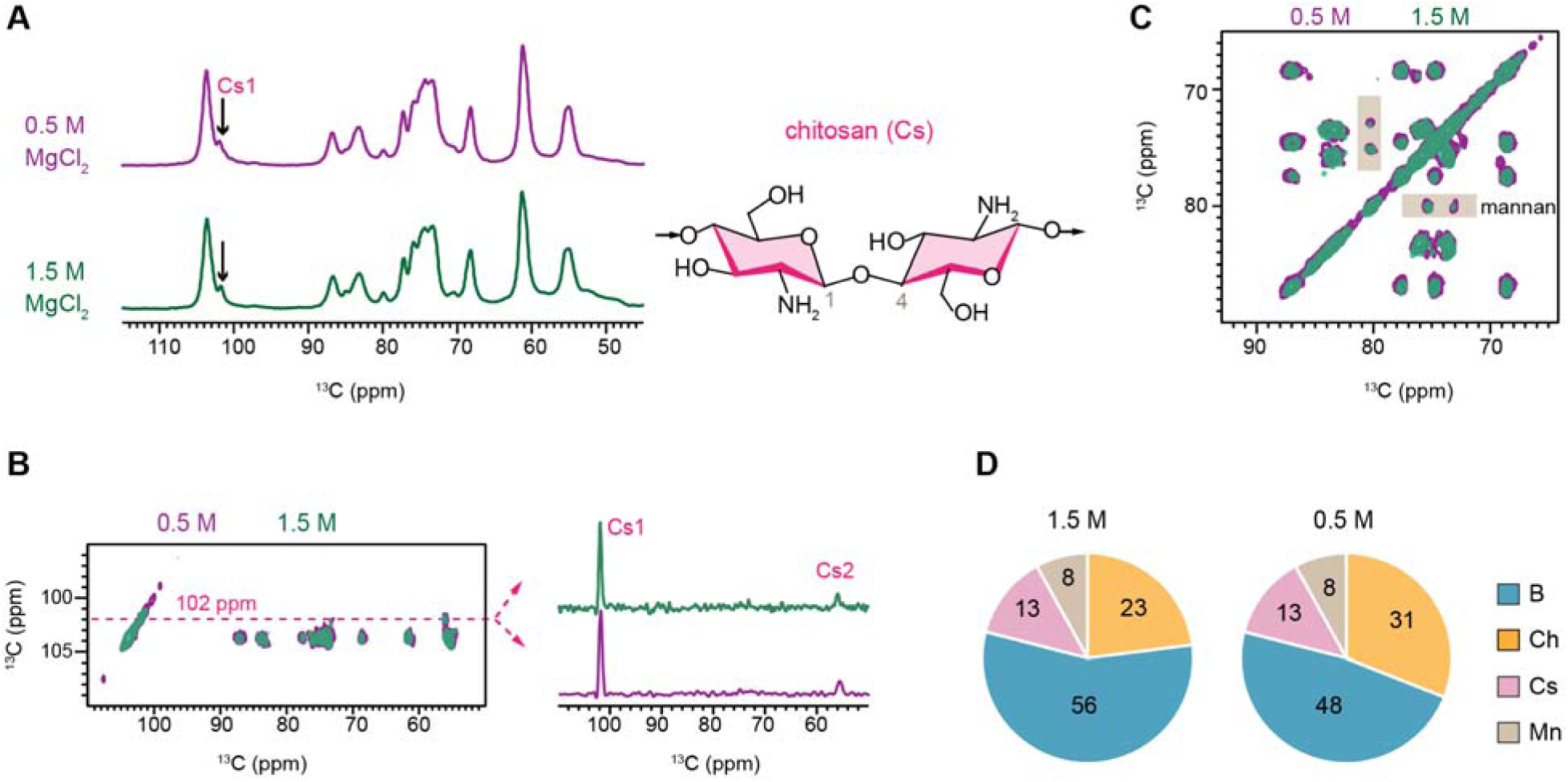
Emergence of chitosan and mannan in *A. atacamensis* under different MgCl_2_ conditions. (**A**) 1D ^13^C CP spectra of 10-day-old *A. atacamensis* samples grown in different MgCl_2_ conditions of 0.5 M (purple; stressful condition) and 1.5 M (green; optimal conditions). The representative structure of chitosan is shown on the side. (**B**) 1D cross-section extracted at 102 ppm showing chitosan signals in both samples. (**C**) Overlay of two 2D ^13^C-^13^C spectra measured on the cultures with 0.5 M and 1.5 M MgCl_2_ concentrations showing mannan signals in both samples. (**D**) Molar fractional composition of the rigid components in *A. atacamensis* samples with 0.5 M and 1.5 M MgCl_2_.

Intensity analysis revealed consistent fractions of chitosan and mannan in the rigid structural domain of the cell wall, accounting for 13% and 8%, respectively, under both MgCl_2_ conditions (**Fig. 2D**). However, the 1.5 M MgCl_2_ condition displayed a higher content of chitin, but lower β-1,3-glucan levels compared to the 0.5 M MgCl_2_ condition. Despite this variation, the combined abundance of these two molecules remained constant across samples, indicating that the rigid cell wall undergoes structural adjustments to adapt to the differing MgCl_2_ concentrations.

The emergence of chitosan and mannan in the rigid domain of the fungal cell wall was determined to be a specific response to MgCl_2_ exposure. This conclusion was drawn by comparing 10-day cultures grown under optimal conditions for MgCl_2_ (1.5 M) and NaCl (1.7 M). The chitosan and mannan signals were exclusively detected in the MgCl_2_ samples, whereas α-1,3-glucan was only observed in the NaCl samples (**Fig. 3A**). These findings confirm that these molecules are uniquely incorporated into the rigid structural phase of the cell wall in response to the specific ionic environment provided by each salt.

**Figure 3.**
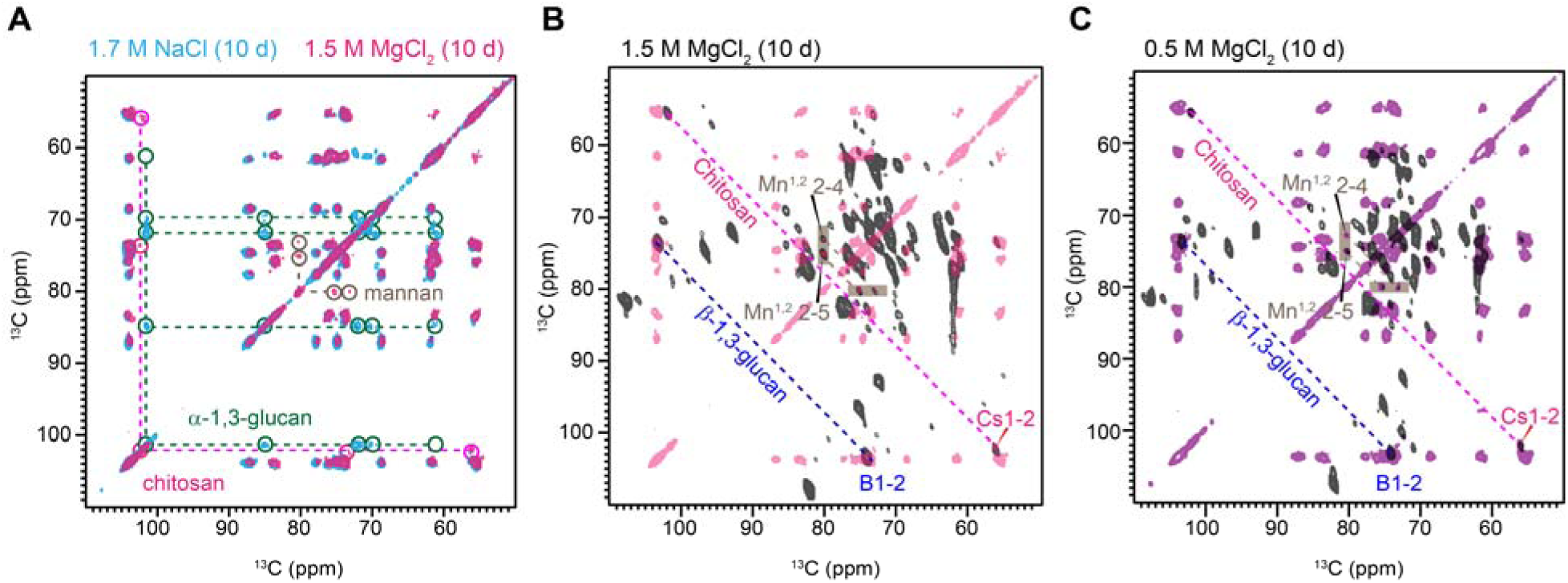
Structural variations in the rigid core of *A. atacamensis*. (**A**) Overlay of two 2D ^13^C-^13^C CORD spectra of *A. atacamensis* samples grown under optimal NaCl and MgCl_2_ conditions for 10 days. The green dashed line and circle highlight α-glucan signals unique to NaCl cultures, while the brown circles indicate mannose signals observed only in MgCl_2_ cultures. Magenta circles highlight chitosan signals found in MgCl_2_ cultures. (**B**) Overlay of ^13^C CORD spectra (pink) with sheared INADEQUATE spectra (black) of the 10-day-old *A. atacamensis* sample cultured under the optimal 1.5 M MgCl_2_ concentration, confirming the identity of mannan peaks. (**C**) Comparison of CORD spectrum (purple) and sheared INADEQUATE spectrum for the stressed 10-day-old sample grown under the sub-optimal 0.5 M MgCl_2_ concentration.

The presence of mannose signals in the rigid regions was confirmed by overlaying the CORD spectra, which selectively detect signals from rigid molecules. These spectra revealed the emergence of mannan signals in the presence of MgCl_2_. Additionally, the sheared DP J-INADEQUATE spectra, which identify mobile components, demonstrated that mannan is a predominant component in the mobile phase (**Fig. 3B, C**). Consistent detection of signals corresponding to mannan C1, C2, and other carbon sites in both spectra indicates the distribution of mannan across both the rigid and mobile fractions in magnesium-saline cultures, irrespective of MgCl_2_ concentration.

### Modulation of GAG level and galactomannan branching in the dynamic phase

The mobile components of *A. atacamensis* predominantly consist of β-1,3-glucan, galactomannan, and galactosaminogalactan (GAG), with their composition varying based on NaCl and MgCl_2_ concentrations (**Fig. 4A**). In prolonged 10-day-old cultures under optimal NaCl conditions (1.7 M), prominent signals corresponding to GAG components, particularly GalN and GalNAc residues, were observed (**Fig. 4B**). These signals were absent in 7-day-old cultures under the same NaCl conditions and in hyperosmotic NaCl cultures with durations of 1-4 days (**Fig. S1)**. Similarly, the GAG signals were detected in 10-day-old cultures under optimal MgCl_2_ conditions (1.5 M) but were absent under stressed conditions (0.5 M). These findings indicate that GAG production is significantly elevated in prolonged cultures under optimal NaCl and MgCl_2_ concentrations, while stressed conditions suppress its synthesis. The elevated content of the anionic exopolysaccharide GAG in mature *A. atacamensis* mycelia may facilitate adhesion to external surfaces, promoting biofilm formation in advanced stages of culture development. This feature correlates with the site of isolation of this strain, as it was found associated with the mucilaginous wall of a cave, suggesting a natural propensity for surface colonization in hypersaline environments(Martinelli et al., 2017).

**Figure 4.**
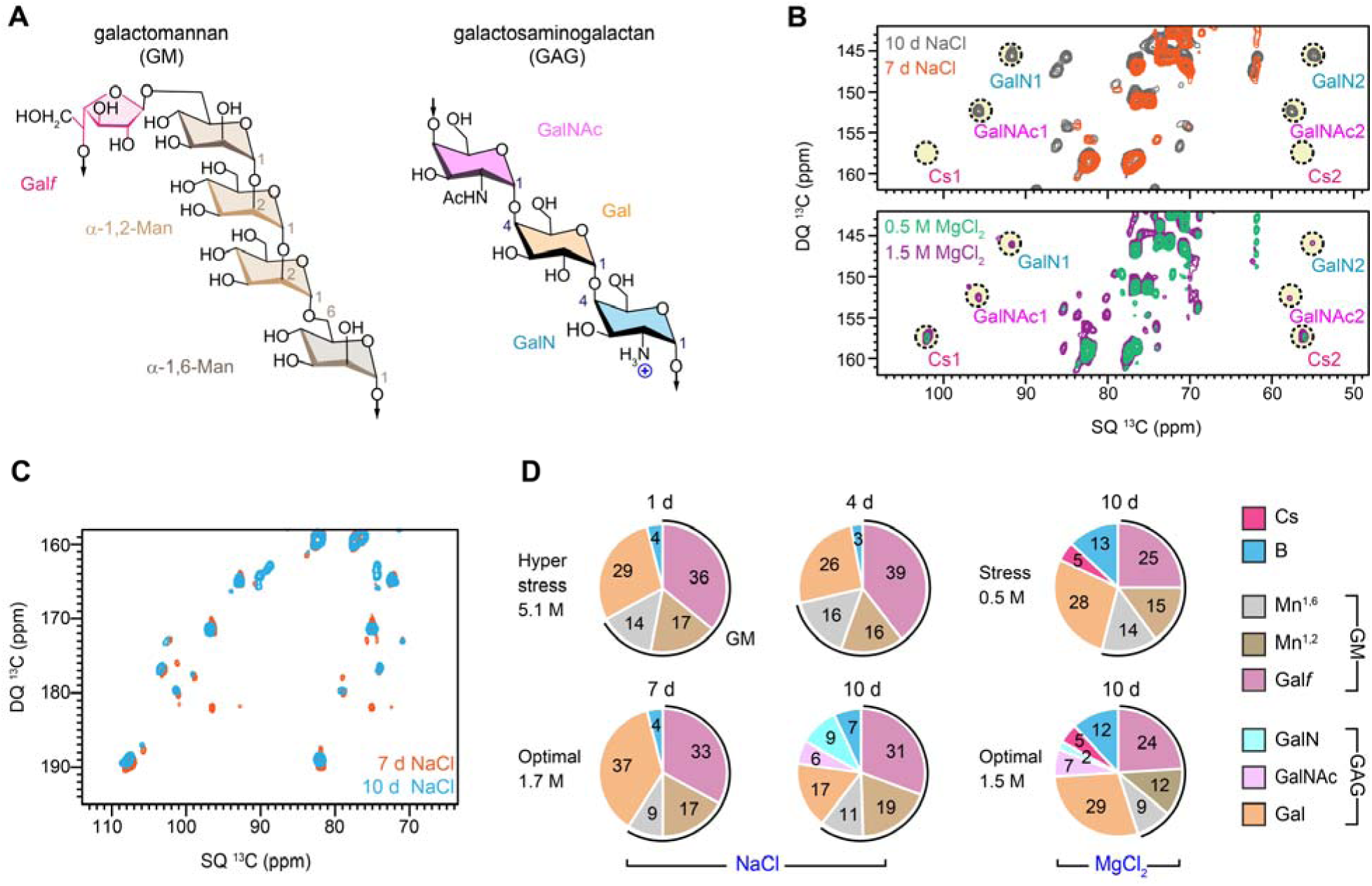
Modifications of mobile carbohydrates of *A. atacamensis*. (**A**) Simplified representation of exopolysaccharide structures of galactomannan and galactosaminogalactan. (**B**) Overlay of 2D ^13^C-^13^C DP J-INADEQUATE spectra of cultures grown under varied saline conditions: 7-day-old (orange) and 10-day-old (grey) samples cultured in NaCl at the optimal condition (1.7M), as well as 10-day-old samples cultured with MgCl_2_ at stressful (0.5 M, green) and optimal (1.5 M, purple) conditions. The spectra reveal the appearance of GAG components with prolonged growth condition in NaCl exposure, while GAG monomers – GalN, GalNAc are prominent under MgCl_2_ conditions, as highlighted by the dashed circles. (**C**) Overlay of C1-C2 region of 2D-DP J INADEQUATE spectra of 7-day-old (orange) and 10-day-old (cyan) samples under optimal NaCl conditions, showing largely consistent signals. (**D**) Molar composition of mobile carbohydrates estimated from resolved signals of DP J-INADEQUATE spectra. The quantity of GM is highlighted using black partial circles.

Signals corresponding to β-1,3-glucan and galactomannan, including its linear α-1,2- and α-1,6-linked backbone and galactofuran sidechain formed by Gal*f* residues, were consistently detected in 7- and 10-day-old cultures under optimal NaCl conditions (**Fig. 4C**), as well as in 1-4-day-old cultures under stressed NaCl conditions. The persistent presence of GM highlights its more consistent role in forming the mobile phase of the cell wall compared to GAG. The branching ratio between the galactomannan sidechain and backbone was remarkably consistent across the four NaCl-treated samples, ranging from 1.16 to 1.27, suggesting that NaCl concentration does not significantly influence galactomannan structure (**Fig. 4D**). In contrast, this branching ratio dropped significantly in MgCl_2_-treated cultures, particularly under stressed conditions, where the ratio decreased to 0.86, indicating a substantially less-branched galactomannan structure.

### MgCl_2_ induces minimal perturbations to cell wall thickness and hydration compared to NaCl

TEM images revealed ultrastructural variations in the cell walls of *A. atacamensis* under different saline conditions. Under hyperosmotic NaCl conditions (5.13 M), the 1-4-day-old cell wall exhibited reduced average thicknesses of 66 nm and 77 nm (**Fig. 5A**, **Table S1**). The 7-10-day-old cell walls grown under optimal NaCl concentrations (1.7 M) or varying MgCl_2_ concentrations displayed increased thickness. Notably, *A. atacamensis* exhibited pronounced cell wall thickening under elevated MgCl_2_ concentrations: Cells grown in optimal MgCl_2_ conditions (1.5 M) demonstrated significantly thicker cell walls (121 nm), whereas those exposed to a more stressful environment with 0.5 M MgCl_2_ had thinner cell walls (99 nm). These findings highlight distinct stress response mechanisms in *A. atacamensis* under magnesium stress, contrasting with the responses observed in *A. sydowii*, where high NaCl concentrations (2.0 M) lead to thicker cell walls compared to optimal NaCl conditions (0.5-1.0 M) and the absence of salt, which acts as a stress condition (Liyanage D Fernando et al., 2023). This structural adaptation appears to be a strategy to counteract osmotic stress and preserve cell integrity in hypersaline environments. A similar response has been observed in the obligate halophilic basidiomycete *W. ichthyophaga*, which undergoes substantial cell wall thickening under high salinity, reaching 0.8 ± 0.2 μm. This response contrasts with related species such as *W. muriae* and *Wallemia sebi*, which exhibit only slight thickening under hypersaline conditions (Kralj Kuncic et al., 2010; J. Zajc et al., 2014; Janja Zajc et al., 2013). The ability of *W. ichthyophaga* to thrive in saturating NaCl concentrations further reinforces the role of structural remodeling as a crucial adaptation to extreme salinity. This evidence raises important questions about how different salts trigger distinct stress responses. Mechanisms in halophilic fungi, particularly in relation to the contrasting effects of kosmotropic versus chaotropic salts on cell wall architecture and remodeling, are also of interest.

**Figure 5.**
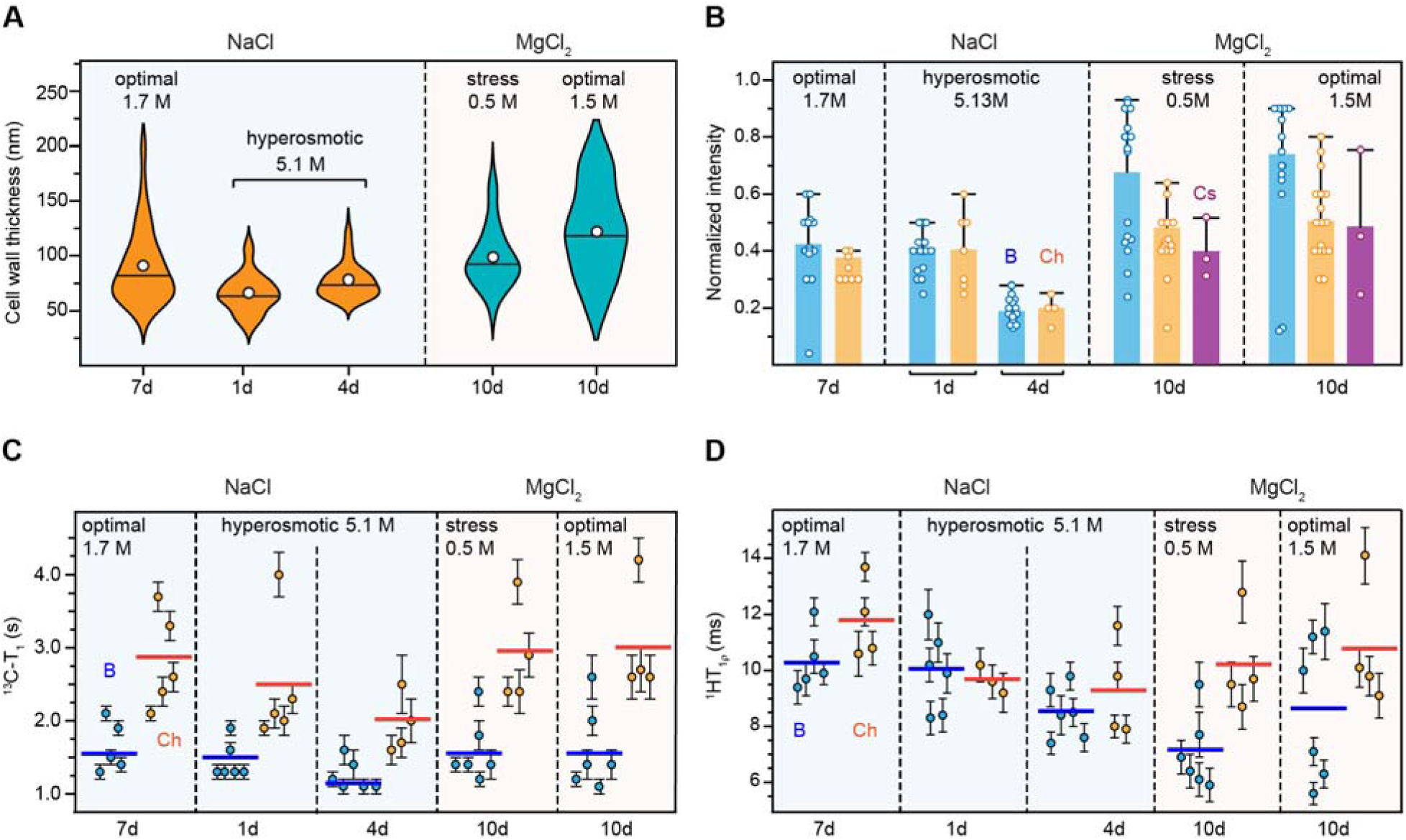
Aging extremophilic fungi develop thickened and rigid cell walls to withstand stress. (**A**) Cell wall thickness of *A. atacamensis* samples cultured under different salt conditions. NaCl cultures are shown in orange (n=120 for all time points), while MgCl_2_ cultures are shown in blue (n= 105 for stress at 0.5M concentration and n=120 for optimal at 1.5M. Open circles represent the mean values, and the horizontal lines indicate the median. (**B**) Normalized water-edited S/S_0_ intensity ratios reflecting the water accessibility of different carbohydrates in the *A. atacamensis* cell wall. Bars represent the mean, and error bars indicate 1.5 × the interquartile range (IQR). Open circles show individual S/S_0_ data points across different carbon sites. Sample sizes: β-1,3-glucan (B: blue, n = 25, 22, 27, 19, 17), chitin (Ch: yellow, n = 7, 6, 9, 20, 23), and chitosan (Cs: purple, n = 3 per MgCl_2_ condition). (**C**) ^13^C-T_1_ relaxation time constants of different carbon sites in chitin (orange) and β-1,3-glucan (blue) in different *A. atacamensis* samples. (**D**) ^1^H-T_1_ relaxation time constants of chitin (orange) and β-1,3-glucan (blue). Error bars represent the s.d. of the fit parameters. Horizontal bars represent the average for each carbohydrate. n values for ^13^C-T_1_ (B: n = 6 for MgCl_2_ conditions, n = 5 for NaCl optimal, n = 6 for NaCl hyperosmotic; Ch: n = 4, 5, 5, 4), and ^1^H-T_1_ (B: n = 6, 5, 6, 6; Ch: n = 4, 4, 3, 4).

These contrasting patterns are consistent with our recent work in *A. sydowii*, where growth under extreme MgCl_2_ concentrations produced transcriptomic profiles that differed sharply from those observed under NaCl, including marked changes in cell wall-related gene expression (González-Abradelo et al., 2025). Thus, MgCl_2_ triggered transcriptional signatures in *A. sydowii* that diverged markedly from those induced by NaCl, revealing ion-specific regulatory programs.

Together, these observations suggest that halophilic fungi can modulate both their wall architecture and gene expression landscapes in a salt-dependent manner, deploying distinct structural and molecular strategies in response to kosmotropic versus chaotropic environments. Such ion-specific remodeling strategies not only deepen our understanding of microeukaryotic biology under extreme salinity but also suggest the possibility of engineering fungal cell walls with tunable properties such as hydrophobicity, porosity, surface charge, and rigidity. This salt-driven customization could enable ad hoc biotechnological applications, including adsorption of organic and inorganic contaminants, enzyme immobilization, and the development of bio-based filtration materials for harsh environments.

In contrast, halotolerant fungi such as *Aspergillus flavus* and *Penicillium roquefortii* exhibit reduced cell wall thickness under 8% NaCl stress, a response that is compensated by increased membrane thickness (Kralj Kunc ic et al., 2010). Similarly, in *Aureobasidium pullulans*, cell wall thickness decreases from 0.20 μm under non-saline conditions to 0.15 μm at 17% NaCl, suggesting an alternative adaptation strategy. The role of osmotic stress in modulating cell wall biogenesis is further emphasized by high glucose concentrations, which have been shown to reduce cell wall biosynthesis in osmosensitive and osmophilic species such as the halophilic species *W. ichthyophaga* and *W. sebi* (Abu-Seidah, 2007). These variations in response have been linked to shifts in molecular composition, particularly changes in β-1,3-glucan and chitin levels, highlighting the diversity of fungal adaptations to osmotic stress influenced by species-specific traits, solute type, and environmental conditions (Kapteyn et al., 1999).

The cell wall of *A. atacamensis* undergoes significant alterations in water accessibility and glucan dynamics under varying saline conditions. The water retention properties of polysaccharides were assessed using the S/S_0_ ratio (**Fig. 5B**), derived from the intensities of the 2D ^13^C-^13^C water-edited spectrum (S, detecting only well-hydrated carbohydrates in the rigid phase) and the control spectrum (S_0_, detecting all rigid carbohydrates). The 4-day-old culture under 5.13 M NaCl exhibited the lowest hydration level, as indicated by the low S/S_0_ values of 0.21-0.22 for both glucans and chitin. This suggests that under hyperosmotic conditions, the fungal cell wall undergoes dehydration, potentially through increased molecular packing and reduced water retention within the polysaccharide matrix. Such structural modifications may help to lower water activity and mitigate osmotic stress by restricting excessive water efflux.

Under hyperosmotic stress, β-1,3-glucan and chitin exhibited nearly identical S/S_0_ ratios within each sample, indicating a uniform hydration level across the rigid core of the cell wall. This finding is unusual, as β-1,3-glucan is typically much more hydrated than chitin due to the natural exclusion of water molecules during chitin microfibril assembly. This trend, particularly observed in other fungal species, including *Aspergillus fumigatus*, *A. sydowii*, *Aspergillus nidulans*, and *Candida* species (Bishoyi et al., 2025; Chakraborty et al., 2021; M.C. Dickwella Widanage et al., 2025; Liyanage D Fernando et al., 2023; Isha Gautam et al., 2025), as well as in *A. atacamensis* grown under optimal NaCl or MgCl_2_ conditions, contrasts with the homogeneous hydration observed in the hyperosmotic samples.

Three possible explanations may account for this atypical hydration pattern: i) the exceptionally thin cell walls produced under hyperosmotic conditions may prevent the formation of hydration gradients, as there is less structural depth to accommodate differential hydration levels; ii) the degree of aggregation and crystallinity of chitin microfibrils under hyperosmotic stress may be reduced, limiting the formation of large crystalline domains that typically exclude water molecules and create localized hydration heterogeneity; and iii) hyperosmotic stress might trigger modifications in carbohydrate crosslinking, altering the structural organization of the polysaccharide network and homogenizing hydration levels across the cell wall matrix.

From an evolutionary and osmoadaptive perspective, this phenomenon may reflect a unique strategy in obligate halophilic fungi, enabling them to maintain cell wall flexibility and mechanical stability under extreme salinity. In contrast, fungi not adapted to high salinity typically exhibit pronounced hydration differences between β-1,3-glucan and chitin, potentially making their cell walls more vulnerable to osmotic fluctuations. This structural plasticity may be a defining feature of fungal haloadaptation, distinguishing obligate halophiles from their halotolerant and non-halophilic counterparts.

Molecules in cultures grown in the presence of MgCl_2_ were significantly better hydrated compared to those in NaCl. The average S/S_0_ ratios were 0.68 and 0.74 under stressed and optimal MgCl_2_ conditions, respectively, markedly higher than the 0.21-0.43 range observed across all NaCl cultures (**Fig. 5B**). Additionally, within each MgCl_2_ sample, the S/S_0_ ratio of β-1,3-glucan was substantially higher than that of chitin/chitosan, indicating that β-1,3-glucan forms the matrix regulating water accessibility, while chitin and chitosan aggregate in the mechanical core. This structural arrangement aligns with the widely observed pattern in many *Aspergillus* species and other species (Chakraborty et al., 2021; Cheng et al., 2024; Malitha C Dickwella Widanage et al., 2024; M.C. Dickwella Widanage et al., 2025; Liyanage D Fernando et al., 2023). These findings suggest that MgCl_2_ exerts a different influence on cell wall hydration than NaCl, allowing for greater water retention within the β-1,3-glucan network and preventing the extreme dehydration observed under hyperosmotic NaCl stress. Unlike NaCl, which induces a progressive loss of hydration and structural rigidity, MgCl_2_ promotes a more balanced hydration profile, potentially helping maintain cell wall plasticity. This could be a key adaptation enabling *A. atacamensis* to tolerate chaotropic environments, as enhanced water retention may protect macromolecular structures from destabilization. Furthermore, the ability to maintain a consistent hydration profile under both stress and optimal MgCl_2_ conditions suggests that *A. atacamensis* employs specific osmoadaptive mechanisms to regulate water dynamics in response to chaotropic stress. This feature may distinguish obligate halophiles from other fungi.

The molecular motions of the cell wall polysaccharides were investigated using ^13^C-T_1_ relaxation (**Fig. 5C** and **Fig. S4**) and ^1^H-T_1_ relaxation (**Fig. 5D** and **Fig. S5**), which probe motions characteristic of the nanosecond and microsecond time scales, respectively. The average ^13^C-T_1_ time constants for β-1,3-glucans and chitin in optimal NaCl culture were 1.6 s and 2.8 s, respectively (**Fig. 5C**). Under 5.1 M NaCl hyperosmotic conditions, the average ^13^C-T_1_ of β-1,3-glucan decreased to 1.5 s (1-day-old sample) and 1.25 s (4-day-old sample). In comparison, the average ^13^C-T_1_ of chitin decreased to 2.5 s and 2.0 s for the two samples, respectively. In contrast, the introduction of MgCl_2_ to the growth medium maintained ^13^C-T_1_ time constants comparable to those of cultures grown in optimal 1.7 M NaCl, with values remaining at 1.6 s for β-1,3-glucan and 2.9-3.0 s for chitin. Thus, while hyperosmotic conditions significantly increased the nanosecond dynamics of polysaccharides in the cell wall, no such effect was observed in the presence of MgCl_2_.

Dynamical changes induced by sodium and magnesium were found to differ on the microsecond and nanosecond timescales. In optimal NaCl conditions, β-1,3-glucan and chitin exhibited very long average ^1^H-T_1ρ_ relaxation time constants of 10.3 ms and 11.8 ms, respectively (**Fig. 5D**). These values were significantly lower in all other cultures. The average ^1^H-T_1ρ_ of β-1,3-glucan was 10 ms and 8.5 ms for the two hyperosmotic samples, and even shorter—8.6 ms and 7.1 ms—for MgCl_2_ cultures. Similarly, the average ^1^H-T_1_ of chitin decreased from the optimal NaCl culture value to 9.7 ms and 9.3 ms for the two hyperosmotic samples, and to 10.8 ms and 10.2 ms for MgCl_2_ cultures. The most dramatic changes were observed for β-1,3-glucan in the 4-day-old hyperosmotic culture with 5.1 M NaCl and for chitin in the stressed 0.5 M MgCl_2_ culture. Therefore, molecules become more dynamic under hyperosmotic conditions and in the presence of MgCl_2_.

### Sodium and magnesium: deciphering the dialogue with the fungal cell wall

The cell wall of *A. atacamensis* exhibits remarkable plasticity in response to sodium and magnesium salts, adapting its composition, hydration, and molecular dynamics to survive under extreme osmotic stress. This flexibility is primarily mediated by β-1,3-glucan and chitin/chitosan, two major structural polysaccharides that function as the primary architectural framework of fungal cell walls. β-1,3-glucan, a key component in most fungal species, acts as a bridging molecule between rigid and flexible domains, dynamically responding to environmental stressors to modulate cell wall integrity and function (Fontaine et al., 2000; N.A.R. Gow et al., 2017; J.P. Latgé & Wang, 2022; Valsecchi et al., 2019). Our data show that β-1,3-glucan comprises 50-70% of the rigid structural domains in all *A. atacamensis* cell walls (**Fig. 1E**). However, its hydration profile and molecular mobility are susceptible to salt type and concentration.

Under hyperosmotic NaCl stress (5.13 M, 4 days), the cell wall undergoes significant dehydration, as evidenced by the marked reduction in water retention capacity (**Fig. 5B**). This dehydration correlates with an increase in both rapid local motions on the nanosecond timescale (**Fig. 5C**) and collective microsecond-scale movements (**Fig. 5D**) for β-1,3-glucan and chitin, suggesting an overall increase in molecular flexibility. The restricted water availability under NaCl stress likely serves as a mechanism to counteract osmotic pressure, ensuring cellular integrity in a hypersaline environment. In contrast, exposure to MgCl_2_ results in a fundamentally different response, in which β-1,3-glucan and chitin/chitosan retain higher hydration levels, promoting a more plastic, hydrated matrix (**Fig. 2E**).

Chitin, a mechanical scaffold in fungal cell walls, forms semi-crystalline fibrils predominantly arranged in an antiparallel configuration (L. D. Fernando et al., 2021; Muszkieta et al., 2014). In all cultures, chitin/chitosan accounts for 30-45% of the rigid cell wall fractions. However, in MgCl_2_-grown cultures, chitin and chitosan coexist at approximately a 2:1 molar ratio. Exposure to MgCl_2_ induces a distinct structural shift, leading to the deacetylation of approximately one-third of the chitin to chitosan (**Fig. 2D**). This transition suggests that MgCl_2_ promotes a more flexible and dynamic cell wall architecture. This feature may contribute to *A. atacamensis*’ ability to withstand chaotropic stress. Additionally, both chitin and β-1,3-glucan exhibit increased molecular mobility under MgCl_2_ stress, further reinforcing the plastic nature of the fungal cell wall across different salt environments.

The production of exopolysaccharides also plays a crucial role in fungal adaptation to saline stress. In *A. atacamensis*, GAG synthesis is upregulated in 10-day-old cultures grown under optimal NaCl or MgCl_2_ concentrations (**Fig. 4B, D**), potentially enhancing biofilm formation and promoting cell adhesion for long-term survival. However, under stress conditions, GAG synthesis is reduced, likely as an energy-conservation strategy that prioritizes cellular maintenance over adhesion-related processes. This behavior contrasts with that observed in *A. sydowii*, where GAG levels remain consistently high (27-36% of mobile polysaccharides), increasing proportionally with rising NaCl concentrations and peaking under hypersaline stress (Liyanage D Fernando et al., 2023). The divergence in GAG regulation between these two fungi highlights species-specific strategies for managing osmotic stress.

Beyond GAG, the role of galactomannan (GM) becomes more pronounced in cultures lacking GAG. GM is consistently detected in NaCl-grown cultures, where it constitutes 59-71% of mobile polysaccharides and maintains stable branching ratios (1.16-1.27, **Fig. 4D**). Interestingly, MgCl_2_ exposure reduces GM content to 44-54% and alters its branching ratio, particularly under MgCl_2_ stress conditions (0.5 M), where an exceptionally low branching degree of 0.86 was observed. This suggests that GM regulation serves as a compensatory mechanism, allowing *A. atacamensis* to fine-tune its cell wall composition and flexibility in response to MgCl_2_-induced stress.

The differential response of *A. atacamensis* to NaCl and MgCl_2_ reflects a broader evolutionary adaptation strategy among halophilic fungi. Increased chitin deacetylation to chitosan under MgCl_2_ exposure suggests a mechanism to enhance cell wall flexibility, which may help counteract chaotropic destabilization. Similarly, the strong dehydration response observed under NaCl stress highlights a structural adaptation aimed at reducing water loss and minimizing osmotic damage. These adaptations align with previous findings in other halophilic fungi, such as *A. sydowii*, where cell wall remodeling plays a central role in extreme osmoadaptation(Liyanage D Fernando et al., 2023).

However, while many halotolerant fungi respond to osmotic stress by thickening their cell walls (Kralj Kuncic et al., 2010; J. Zajc et al., 2014; Janja Zajc et al., 2013), *A. atacamensis* appears to adopt a more dynamic approach, adjusting its polysaccharide composition, hydration levels, and molecular organization in a salt-specific manner. This plasticity could provide *A. atacamensis* with a unique advantage over fungi that rely solely on rigidification under salt stress. In non-halophilic fungi, exposure to high salt concentrations typically leads to reduced growth, increased metabolic burden, and structural weakening of the cell wall, underscoring the importance of specialized osmoadaptive traits in true halophiles (Gunde-Cimerman et al., 2018).

A decade after the initial studies on chaotolerance in fungi (J. Zajc et al., 2017), progress in this field remains limited, with many fundamental questions still unanswered. The findings presented here provide key insights into how *A. atacamensis* restructures its cell wall under NaCl and MgCl_2_ stress, yet critical knowledge gaps persist. Future research should focus on elucidating the molecular pathways governing chitin deacetylation under MgCl_2_ stress, as this process appears to be a key adaptive strategy for enhancing cell wall flexibility in chaotropic environments. Additionally, a deeper investigation into the functional role of GM downregulation under MgCl_2_ stress is necessary to understand its implications for cell wall permeability and mechanical properties. Further integrating transcriptomic and proteomic analyses will be essential for establishing direct links between gene expression, enzymatic activity, and structural changes observed in the fungal cell wall. Comparative studies across other halophilic fungi will help determine whether similar plasticity exists across different evolutionary lineages or if *A. atacamensis* represents a uniquely versatile adaptation model. Expanding these efforts will enhance our understanding of fungal extremophiles and may contribute to biotechnological applications, including the development of salt-tolerant fungal strains and biomaterials suitable for extreme environments.

## CONCLUSIONS

This study provides a molecular-level description of how an obligate halophilic fungus restructures its carbohydrate-based cell wall in response to extreme kosmotropic and chaotropic environments. Using intact-cell solid-state NMR, we demonstrate that *A. atacamensis* maintains a β-1,3-glucan-and chitin-dominated rigid core under a wide range of NaCl concentrations. In contrast, exposure to MgCl_2_ induces pronounced remodeling of cell wall carbohydrates, including partial chitin deacetylation to chitosan and rigid-phase incorporation of mannan, accompanied by increased wall thickness, hydration, and polymer mobility. These features suggest that chaotropic stress is mitigated through the formation of a more hydrated and dynamic polysaccharide network, in sharp contrast to the dehydration-driven response observed under hyperosmotic NaCl stress. Together, these results reveal that *A. atacamensis* employs salt-specific carbohydrate strategies to preserve cell wall integrity under extreme conditions, highlighting the importance of polymer chemistry, hydration, and dynamics in fungal halophily and chaotolerance. This work expands current understanding of fungal cell wall adaptation beyond halotolerance and establishes a framework for exploring carbohydrate-based survival strategies in extremophilic eukaryotes, with potential implications for the design of salt-tolerant biomaterials and fungal biotechnology.

## SUPPORTING INFORMATION

Supporting Information: Figures S1-S6, Tables S1-S7, and Supplementary Methods for additional solid-state NMR spectra and transcriptomic analysis.

## ACKNOWLEDGMENT

This work was supported by the National Institutes of Health (NIH) grant R01AI173270 to T.W. This work was also supported by the RYC2022-037554-I project funded by MCIN/AEI/10.13039/501100011033 and FSE+ (RAB-G). The Secretary of Science, Humanities, Technology and Innovation (SECIHTI) of the Government of Mexico supported this work: Projects 1059, 315114, and SEP-CB-285816 (RAB-G). This study was also supported by funding from the Slovenian Research Agency to Infrastructural Centre Mycosmo (MRIC UL, I0-0022), programs P4-0432 and P1-0198 (NG-C). This work utilized resources from the New Mexico State University (NMSU), NM, USA High Performance Computing Group, which is directly supported by the National Science Foundation (OAC-2019000), the Student Technology Advisory Committee, and New Mexico State University and benefits from inclusion in various grants (DoD ARO-W911NF1810454; NSF EPSCoR OIA-1757207; Partnership for the Advancement of Cancer Research, supported in part by NCI grants U54 CA132383 (NMSU)). The authors sincerely thank Prof. Adriana Romero-Olivares, PI of the Fungal Ecology Laboratory at NMSU, for her technical support. The authors also thank Dr. Irina Jiménez-Gómez (Universidad Autónoma del Estado de Morelos, Mexico; currently at UTHealth Houston, TX, USA) for her support in establishing the culture protocol for *A. atacamensis* EXF-6660 used in this study. GV-M received a PhD fellowship from SECIHTI, Mexico. Prof. Armando Azua-Bustos is acknowledged for the deposition of the strain *Aspergillus atacamensis* (Holotype Chile: *A. atacamensis* Zalar, Azúa-Bustos, Gunde-Cimerman, sp. nov) in the Culture Collection ExF within IC Mycosmo.

## CRediT Authorship Contribution Statement

**Isha Gautam**: Writing – original draft, Investigation, Formal analysis. **Gisell Valdés Muñoz**: Writing – review & editing, Investigation, Formal analysis. **Aswath Karai**: Writing – review & editing, Investigation. **Jean-Paul Latgé**: Writing – review & editing, Conceptualization. **Yordanis Pérez-Llano**: Investigation. **Nina Gunde-Cimerman**: Writing – review & editing, Resources. **Ramón Alberto Batista-García**: Writing – review & editing, Conceptualization, Funding acquisition. **Tuo Wang**: Writing – review & editing, Conceptualization, Funding acquisition.

## Abbreviations

CORD: COmbined R2^n^ν-Driven
CP: cross polarization
CryoEM: cryo-electron microscopy
DP: direct polarization
GAG: galactosaminogalactan
GM: galactomannan
INADEQUATE: Incredible Natural Abundance DoublE QUAntum Transfer Experiment
MAS: magic-angle spinning
TEM: Transmission electron microscopy

