## Supplementary Material for "Chaotropic Ions Reshape the Cell Wall of the Obligate Halophile *Aspergillus atacamensis*: Insight from Solid-State NMR"

^4^ Department of Biology, Biotechnical Faculty. University of Ljubljana. Ljubljana, Slovenia

* Correspondence and requests for materials should be addressed to T.W.

 and RAB-G

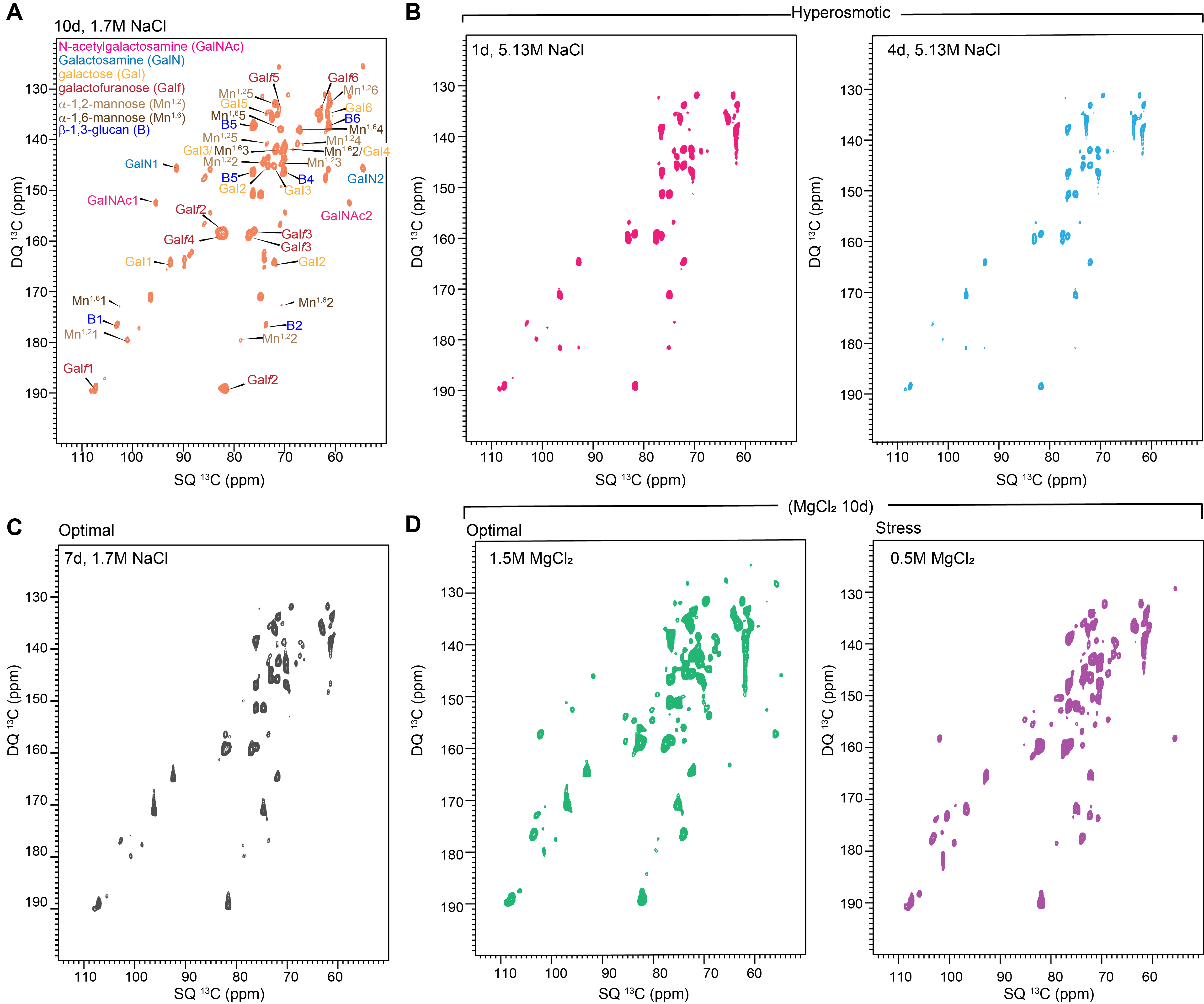

**Figure S1**. **Changes in mobile carbohydrates under high salinity and maturity**. (**A**) 2D ^13^C DP *J*-INADEQUATE spectra of A. atacamensis samples under optimal conditions (1.7 m NaCl) for 7 days, with all glucans assigned and color-coded. (**B**) 2D ¹³C DP *J*-INADEQUATE spectra of samples grown in 5.13 M NaCl for 1 day (pink) and 4 days (blue) under the hyperosmotic conditions. (**C**) Spectrum of the optimal NaCl condition (1.7 M). (**D**) Spectrum of samples grown in 0.5 M (stress, green) and 1.5 M (optimal, purple) MgCl_2_ for 10 days. All spectra were acquired at 800 MHz at 15 kHz MAS.

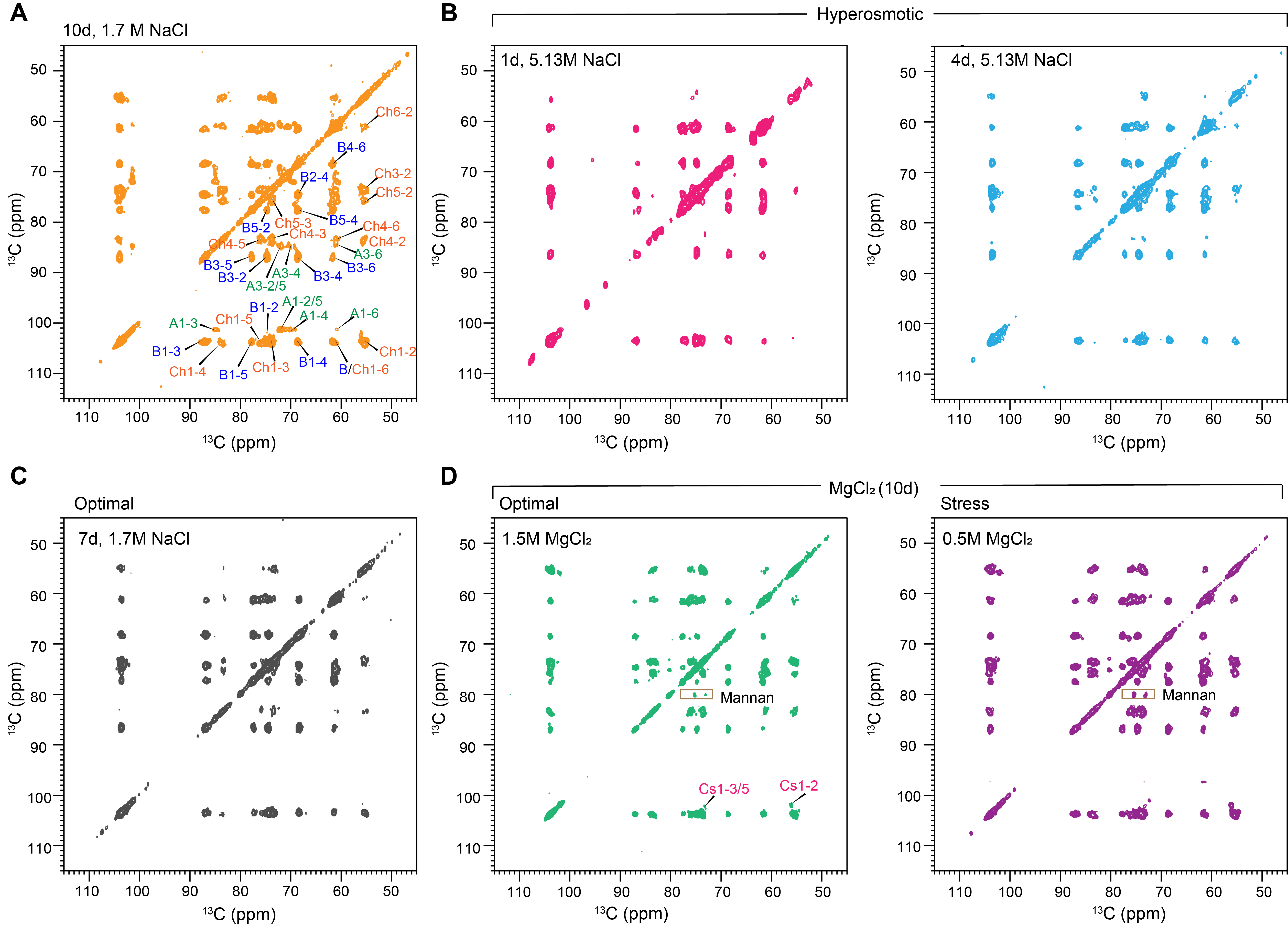

**Figure S2. Changes in rigid carbohydrates with different salinity and culture duration**. (**A**) 2D ¹³C-¹³C CORD spectra of A. atacamensis cultured under optimal conditions for an extended time (10 days) with assigned and color-coded glucans. (**B**) Spectra of samples grown in 5.13 M NaCl for 1 day (pink) and 4 days (blue) under hyperosmotic conditions. (**C**) Optimal 1.7 M NaCl sample grown for 7 days. (**D**) Green and purple spectra represent samples grown in 1.5 M (optimal) and 0.5 M (stress) MgCl_2_, respectively, showing new mannan peaks. All spectra were recorded at 800 MHz at 15 kHz MAS.

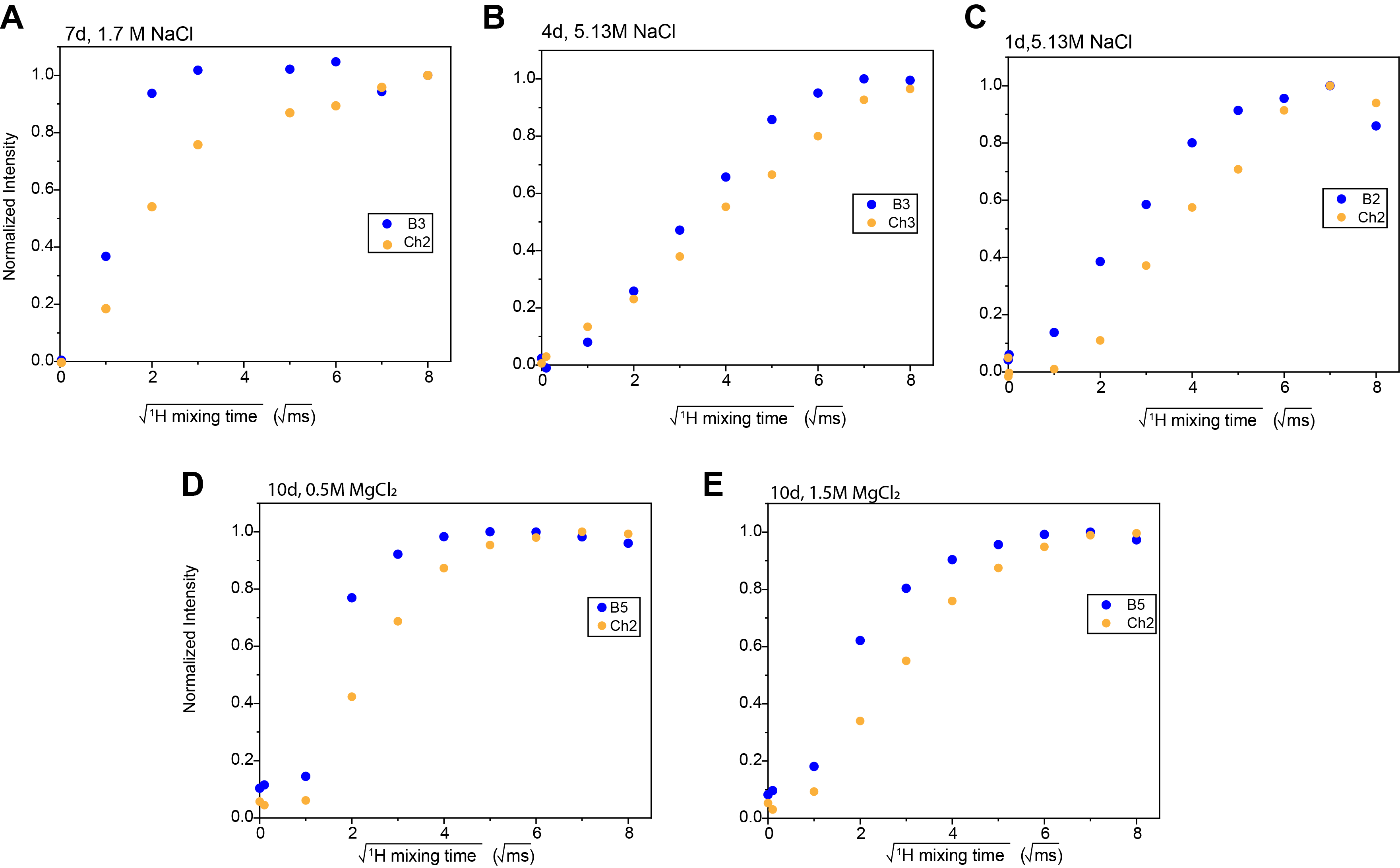

**Figure S3. Water-edited intensity buildup curves for carbohydrates of A. atacamensis**. (**A-C**) Water-edited buildup curves of polysaccharides in A. atacamensis cell walls under optimal conditions (1.7 M NaCl, 7 days) and hyperosmotic conditions (5.13 M NaCl for 1 day and 4 days). The optimal sample (7 days) shows rapid buildup, while hyperosmotic conditions result in slower buildup. (**D**, **E**) Water-edited buildup curves for samples grown in 0.5 M and 1.5 M MgCl_2_, showing enhanced hydration with longer water buildup times in both samples. Data were collected on a 400 MHz (9.4 Tesla) spectrometer at 10 kHz MAS and 277 K. Blue dots represent β-1,3-glucan, orange dots represent chitin, and symbols indicate specific polysaccharide carbons.

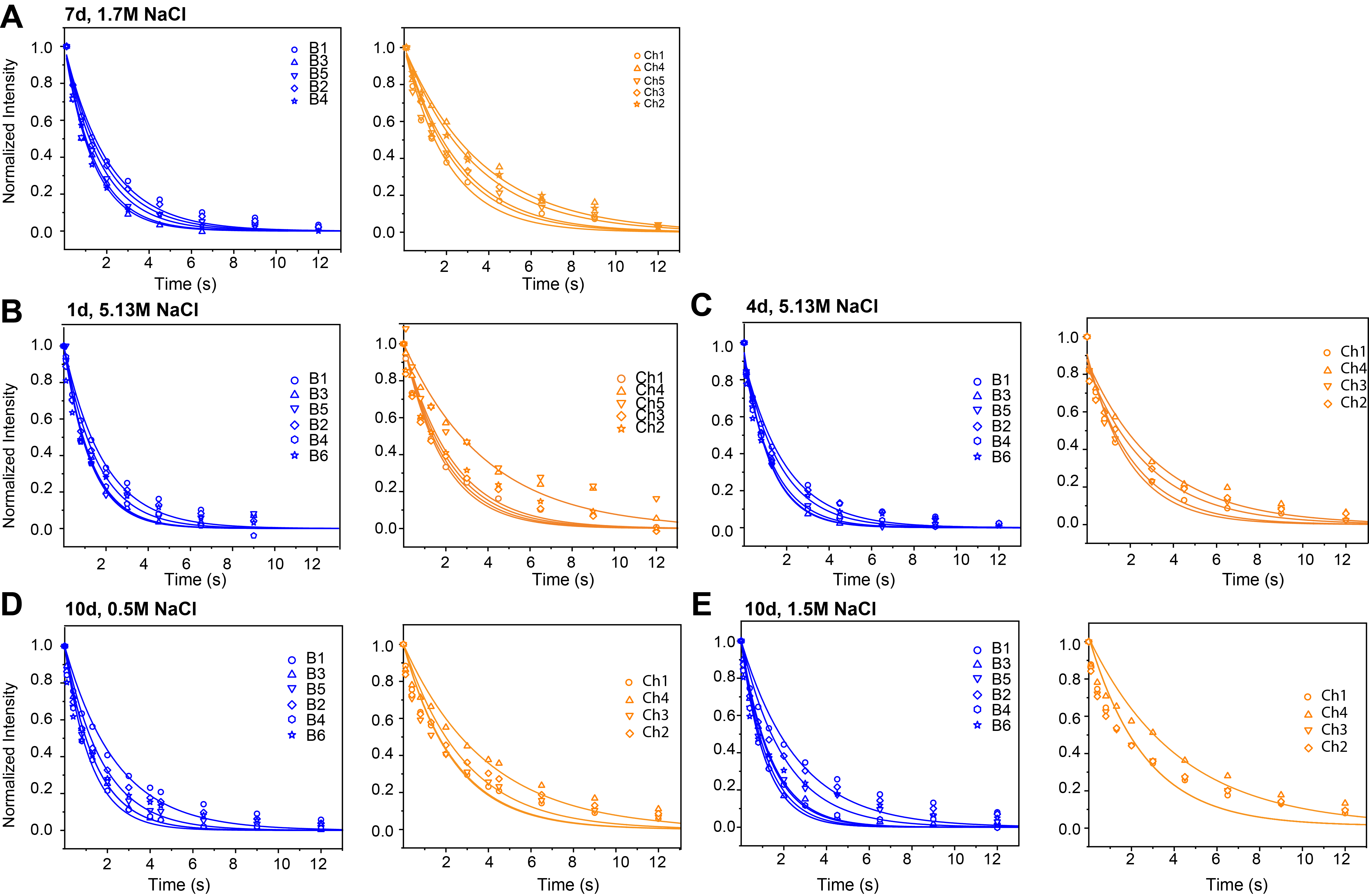

**Figure S4.** **^13^C-T_1_** **NMR relaxation curves of key carbohydrates**. (**A-C**) ^13^C-T_1_ relaxation curves of polysaccharides in A. atacamensis cell walls under optimal conditions (1.7 M NaCl, 7 days) and hyperosmotic conditions (5.13 M NaCl for 1 day and 4 days). (**D**, **E**) Relaxation curves for samples grown in MgCl_2_ at 0.5 M and 1.5 M (optimal). Data were collected on a 400 MHz (9.4 Tesla) spectrometer at 10 kHz MAS, with single-exponential fits. Blue lines represent β-1,3-glucan, orange lines represent chitin, and symbols indicate specific polysaccharide carbons.

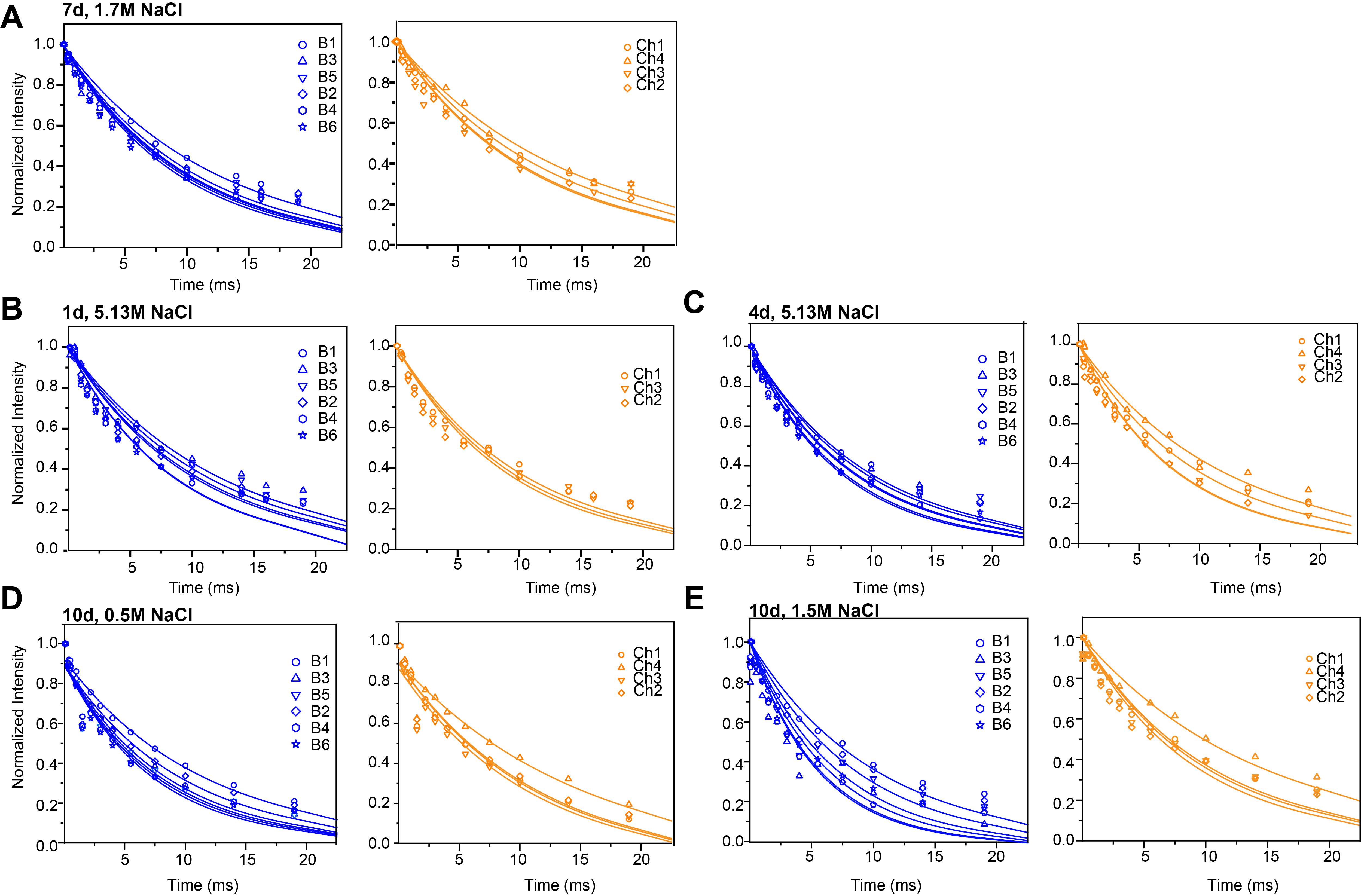

**Figure S5.** **^1^H‒T_1ρ_** **relaxation curves of A. atacamensis** **carbohydrates**. (**A-C**) ^1^H-T_1ρ_ relaxation curves of polysaccharides in cell walls under optimal conditions (1.7 M NaCl, 7 days) and hyperosmotic conditions (5.13 M NaCl for 1 day and 4 days). (**D, E**) Relaxation curves for samples grown in MgCl_2_ at 0.5 M (stress) and 1.5 M (optimal). Data were collected on a 400 MHz (9.4 Tesla) spectrometer at 10 kHz MAS, with single-exponential fits. Blue lines represent β-1,3-glucan, orange lines represent chitin, and symbols indicate specific polysaccharide carbons.

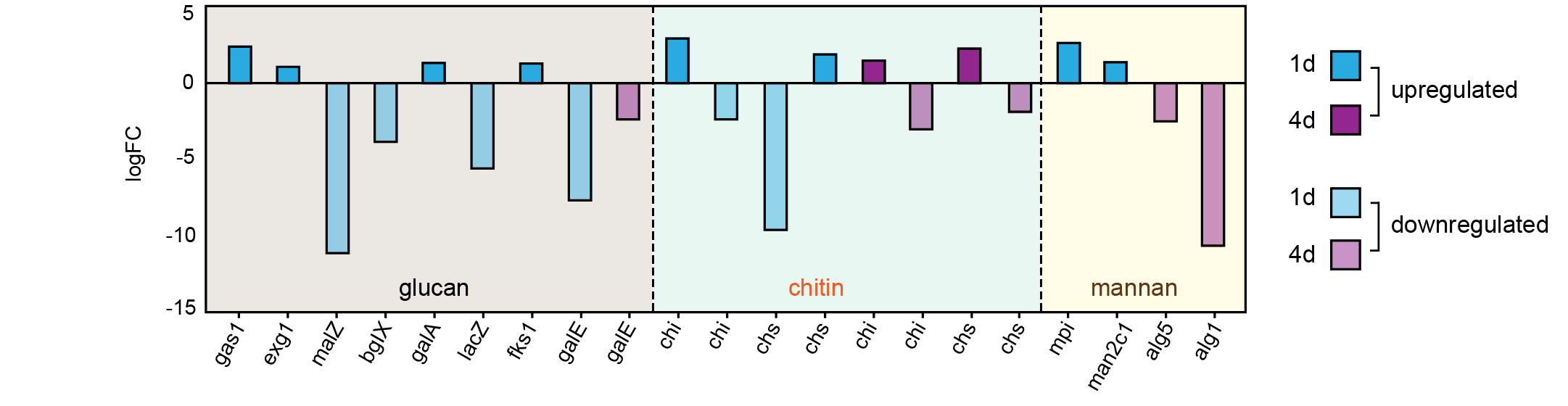

**Figure S6.** **Transcriptomic data on *A. atacamensis* under hyperosmotic stress.** (**A**) Log-fold changes (logFC) of genes involved in the biosynthesis of glucan, chitin, and mannan in *A. atacamensis* at 1 day (blue) and 4 days (purple) under hyperosmotic stress with 5.13 M of NaCl. Genes associated with β-1,3-glucan synthesis, including β-1,3-glucanosyltransferase (*gas1*, logFC=2.4) and β-1,3-glucan synthase (*fks*, logFC=1.47) were upregulated. Genes related to chitin metabolism (*chi* and *chs*) exhibited a downregulation trend

**SUPPLEMENTARY METHODS**

**Transcriptomic analysis.** Fungal mycelium was harvested by centrifugation and cryogenically ground to ensure optimal RNA preservation. Total RNA was extracted using the TRIzol method ^1^. RNA quality was evaluated through capillary electrophoresis using an Agilent Bioanalyzer 2100, and high-quality samples (RIN > 7) were selected for library preparation with the TruSeq Stranded mRNA LT Sample Prep Kit. Sequences were obtained using an Illumina platform with 150-bp paired-end reads, yielding more than 80 million reads per sample.

Following demultiplexing, sequencing read quality was enhanced using Trimmomatic (version 0.39), which filtered low-quality reads, removed 5′ and 3′ adaptors, and removed ^2^, and removed highly overrepresented sequences. Further error correction of Illumina RNA sequencing reads was performed using Rcorrector ^2^. The transcriptome was assembled de novo using Trinity 2.10.0, generating contigs from short-read RNA sequences ^3, 4^. Transcriptome completeness was then assessed using BUSCO 4.0.5, ensuring a high-quality reference for downstream analyses ^5, 6^.

Functional annotations were generated using Blast2GO, employing a blast-based workflow targeting the Fungi section of the nr database. Gene ontology (GO) and InterPro annotations were retrieved through this high-throughput platform ^7^. Transcript abundances were quantified using Kallisto (version 0.46.1) ^8^, while RUVseq was applied to the unnormalized count data to mitigate unwanted variations in RNA sequencing reads ^9^. Differential expression analysis was conducted using the edgeR software package (version 1.29.5) ^10^. GO enrichment testing was performed on the OmicsBox platform ^7^, using Fisher's exact test to control the false discovery rate for GO term enrichment. Finally, pathway enrichment analysis was conducted using KEGG Mapper, with KEGG annotations generated by the GhostKOALA pipeline ^11^.

**Table S1.** **The average cell wall thickness of the extremophile A. atacamensis**. Results are presented as the mean ± standard deviation. The number of individual cells measured is denoted as the n value for each fungal cell sample. Statistical analysis was performed using one-tailed and two tailed assuming unequal variance at 95% confidence level (p <0.05).

| **Sample** | **NaCl** | | | **MgCl_2_** | |
| --- | --- | --- | --- | --- | --- |
|  | **Hyperosmotic (5.17 M)** | | **Optimal (1.7 M)** | **Stressful (0.5 M)** | **Optimal (1.5 M)** |
|  | **1 day** | **4 days** | **7 days** | **10 days** | **10 days** |
| Average cell wall thickness (nm) | 66 ± 18 | 77 ± 16 | 91 ± 35 | 99 ± 29 | 121 ± 43 |
| n | 120 | 120 | 120 | 105 | 120 |

**Table S2. Solid- state NMR experimental parameters for *A. atacamensis.*** The experimental parameters include the ^1^H Larmor frequency, total experiment time (t), recycled delay (d1), number of scans (NS), The number of points for the direct (td2) and indirect (td1) dimensions, the acquisition time of the direct dimension (aq2) and the evolution time of indirect dimension (aq1), mixing time (t_m_), and T filter times (t_z_). * Indicates the water-polysaccharide spin diffusion and the DARR mixing time.

|  | **Experiments** | **T**  **(K)** | **B_0_**  **(T)** | **ν_MAS_**  **(kHz)** | **d1**  **(s)** | **NS** | **td2** | **td1** | **aq2**  **(ms)** | **aq1**  **(ms)** | **t_m_**  **(ms)** | **t_z_** |
| --- | --- | --- | --- | --- | --- | --- | --- | --- | --- | --- | --- | --- |
| 1D | CP | 293 | 18.8 | 15 | 2 | 1024 | 3600 |  | 18 |  |  |  |
|  | DP | 293 | 18.8 | 15 | 2 | 512 | 3600 |  | 18 |  |  |  |
|  |  |  |  |  | 35 |  |  |  |  |  |  |  |
|  | Water-edited | 277 | 9.4 | 10 | 2 | 512 | 2000 |  | 16 |  | 0, 1, 4,9,16,25,36,49,64, 81,100 |  |
|  | 1D Torchia ^13^C-T_1_ | 298 | 9.4 | 10 |  | 512 | 2000 |  | 16 |  |  | (0.1, 0.4,0.8, 1.3,2, 3, 4.5, 6.5, 9, 12) s |
|  | ^1^H T_1ρ_ relaxation | 298 | 9.4 | 10 | 2 | 512 | 1400 |  | 16 |  |  | SL (0.1-19 ms) |
| 2D | 2D CORD | 293 | 18.8 | 15 | 2 | 32 | 3200 | 400 | 16 | 5.2 | 50 |  |
|  | DP J INADEQUATE | 293 | 18.8 | 15 | 2 | 32 | 2800 | 472 | 14 | 5.2 |  |  |
|  | Water-edited control | 277 | 9.4 | 10 | 2 | 64 | 2000 | 220 | 16 | 5.5 | 0/50* | T_2_ 10^-4^ ms |
|  | Water-edited | 277 | 9.4 | 10 | 2 | 64 | 2000 | 220 | 16 | 5.5 | 4/50* | T_2_ 1.2 ms  T_2_ 1.3ms |

**Table S3. ^13^C chemical shift (ppm) of biomolecules in the cell wall in *A. atacamensis*.** The ^13^C chemical shifts reported here are on the TMS scale.

| **Biomolecules** | **C1** | **C2** | **C3** | **C4** | **C5** | **C6** | **Methods** | **Cell wall Portion** | **References** |
| --- | --- | --- | --- | --- | --- | --- | --- | --- | --- |
| β-1,3-glucan | 103.9 | 74.7 | 87.0 | 68.6 | 77.7 | 61.6 | ^13^C-^13^C CORD | Rigid | Shim et al. 2007^12^  Bhanja et al 2014^13^  Dickwella et al 2024^14^  Fontaine et al. 2011^15^ |
| α-1,3-glucan (A) | 101.4 | 72.0 | 84.9 | 70.1 | 72.0 | 60.8 |  |  |  |
| Chitin | 103.9 | 55.1 | 73.7 | 83.4 | 75.9 | 61.0 |  |  |  |
| Chitosan (Cs) | 102.0 | 55.8 | 67.2 | - | - | - |  |  |  |
| α-Mn^1,2^ | 102.4 | 80.1 | 74.1 | - | 75.1 | - |  |  | Kirui et al.2022^16^  Dickwella et al 2024^14^ |
| α-Mn^1,2^ | 101.5 | 79.3 | 71.1 | 68.0 | 72.4 | 61.7 | ^13^C-^13^C  *J*- DP INADEQUATE | Mobile | Latge et al. 1994^17^ |
| α-Mn^1,6^ | 103.1 | 71.1 | 72.5 | 67.4 | 71.4 | 61.6 |  |  | Dickwella et al 2024^14^ |
| Gal*f* | 107.8 | 82.3 | 76.5 | 82.7 | 71.7 | 61.2 |  |  | Chakraborty et al. 2021^18^ |
| Gal | 93.1 | 72.5 | 72.7 | 70.7 | 74.2 | 62.0 |  |  |  |
| GalN | 92.0 | 55.1 | - | - | - | - |  |  | Poulhazan et al. 2021^19^ |
| GalNAc | 96.0 | 57.9 | - | - | - | - |  |  | Fontaine et al. 2011^15, 20^ |

**Table S4: Relative molar composition of the rigid cell of *A. atacamensis*.** The molar composition of rigid components was calculated from the integrals of well-resolved cross peaks of 2D ^13^C-^13^C CORD spectra. Chemical shifts are reported for both single quantum dimensions (SQ), along with the integrated peak volumes for each carbon cross-peak. A dash line (-) indicates signals that were not detected. The molar composition is listed as an average percentage, with the error defined as the standard error.

| **Cross peaks** | **SQ** | **SQ** | **1.17 M NaCl** | | **1.17 M NaCl** | | **5.13 M NaCl** | | **5.13 M NaCl** | | **0.5M MgCl_2_** | | **1.5M MgCl_2_** | |
| --- | --- | --- | --- | --- | --- | --- | --- | --- | --- | --- | --- | --- | --- | --- |
|  |  |  | **10 d** | | **7 d** | | **1 d** | | **4 d** | | **10 d** | | **10 d** | |
|  |  |  | **Abs (10^10)** | **Avg %** | **Abs (10^10)** | **Avg %** | **Abs (10^10)** | **Avg %** | **Abs (10^10)** | **Avg %** | **Abs (10^10)** | **Avg %** | **Abs (10^10)** | **Avg %** |
| B1-3  B1-5  B1-4  B1-2  B3-5  B3-2  B3-4  B3-6  B5-6  B5-2  B5-4  B4-6  B2-4 | 103.8  103.7  103.6  -  87  86.9  87  86.9  -  77.5  77.6  -  74.4 | 87  77.7  68.5  -  77.7  74.6  68.6  61.7  -  74.7  68.5  -  68.5 | 330  130  230  -  180  290  270  100  -  130  270  -  270 | 48±0.4 | 170  110  140  380  120  190  180  85  310  120  200  190  - | 70±0.8 | 40  8.8  30  -  22  46  42  17  -  21  48  47  - | 62±0.5 | 62  28  47  -  39  75  50  29  -  -  69  55  - | 60±0.5 | 200  110  180  -  130  240  250  99  -  130  -  220  - | 31±0.1 | 34  15  20  -  22  35  36  7.3  -  -  -  34  - | 23±01 |
| Ch1-4  Ch1-5  Ch1-2  Ch4-5  Ch4-3  Ch4-6  Ch4-2  Ch5-2  Ch3-2  Ch6-2 | 103.9  104.1  103.3  83.2  83.6  83.3  83.5  75.9  73.5  61.1 | 83.6  76.1  55.4  75.9  73.8  61.2  55.4  55.3  55.5  55.4 | 120  65  350  120  180  -  130  98  290  54 | 34±0.3 | 48  53  120  90  100  46  69  -  120  44 | 30±0.1 | 23  34  -  -  -  -  -  3.4  28  8.9 | 38±0.3 | 38  58  33  25  14  49  14  -  -  - | 40±0.3 | 220  -  440  290  350  180  210  170  440  110 | 48±0.5 | 61  -  110  60  87  29  46  24  120  31 | 56±06 |
| Mn2-5  Mn2-3 | 80  80 | 75.3  73 | -  - |  |  |  | -  - |  | -  - |  | 60  30 | 8±0 | 11  6 | 8±0 |
| Cs1-2  Cs1-3 | 102.6  102.4 | 55.6  73.5 | 36  29 | 7±0 | -  - |  | -  - |  | -  - |  | 93  58 | 13±0 | 20  10 | 13±0.1 |
| A1-3  A1-2  A1-5  A1-4  A1-6  A3-5  A3-2  A3-4 | 101.3  101.1  101.2  101  101.2  84.7  84.4  84.6 | 84.8  71.9  70  69.1  60.8  72  70.3  69.2 | 59.9  140  23  13.6  13.9  85  35  14 | 11±0 | -  -  -  -  -  -  -  - | -  -  -  -  -  -  -  - | -  -  -  -  -  -  -  - |  | -  -  -  -  -  -  -  - |  | -  -  -  -  -  -  -  - |  | -  -  -  -  -  -  -  - |  |

**Table S5. Relative molar composition of the mobile cell wall of *A. atacamensis*.** The Molar composition of mobile components was determined from the integrals of well-resolved signals in 2D ^13^C-^13^C J- INADEQUATE spectra. Chemical shifts are provided for both the single-quantum (SQ) and double-quantum (DQ) dimensions, along with the integrated peak volumes for each carbon spin pair. A dash line (-) indicates signals that were not detected. The molar composition is expressed as the average percentage, with the error defined as the standard errors.

| Biomolecules | **Carbon sites** | **5.13 M NaCl** | | | | **5.13 M NaCl** | | **1.7 M** | | **1.7 M** | | **0.5 M MgCl_2_** | | **1.5 M MgCl_2_** | |
| --- | --- | --- | --- | --- | --- | --- | --- | --- | --- | --- | --- | --- | --- | --- | --- |
|  |  | **1 d** | | | | **4 d** | | **7 d** | | **10 d** | | **10 d** | | **10 d** | |
| Gal*f* |  | **SQ** | **DQ** | **Abs (10^10)** | **Avg %** | **Abs (10^10** | **Avg %** | **Abs (10^10)** | **Avg %** | **Abs (10^10** | **Avg %** | **Abs (10^10** | **Avg %** | **Abs (10^10** | **Avg %** |
|  | C1  C2  C2  C3  C3  C4  C5  C6 | 108.3  81.9  82  77.7  77.7  83.2  73  63.5 | 188.8  188.8  158.8  158.4  160.2  159.6  136.5  136.1 | 4600  4300  2000  3700  2100  3900  5000  720 | 36±0.2 | 4200  2800  2700  2600  2300  3000  1300  1400 | 39±0.1 | 4300  3800  7300  7300  320  400  460  920 | 33±0.2 | 2000  3000  6000  400  0  0  3000  4000 | 31±0.2 | 1000  10000  2000  2000  7000  500  2000  7000 | 25±0.1 | 4000  4000  5000  5000  700  400  800  600 | 24±0.1 |
| Mn^1,2^ | C1  C2  C2  C3  C3  C4  C4  C5  C5  C6 | 101.1  79  79.1  70.4  71.1  67.6  67.8  73.6  72.2  61.7 | 179.7  179.8  149.6  150.1  138.8  138.9  141.2  141.3  133.6  133.4 | 3810  398  2050  268  -  -  -  -  870  2000 | 17±0.07 | 2900  130  140  810  -  -  -  -  1400  3100 | 16±0.06 | 2900  4300  3800  320  630  590  330  190  -  - | 17±0.07 | 4300  5000  300  130  1200  1300  -  -  770  2300 | 19±1 | 4300  5000  3000  130  1200  1300  -  -  770  2300 | 15±0.05 | 3000  2300  2200  860  -  -  280  160  1600  930 | 12±0.04 |
| Mn^1,6^ | C1  C2  C2  C3  C3  C4 | 102.4  71  70.9  74.3  74.2  68.6 | 172.8  172.6  144.9  144.7  142.7  142.8 | 3090  3330  116  124  339  687 | 14±0.06 | 3500  1100  0  0  519  499 | 16±0.07 | 100  820  2000  780  -  - | 9±0.03 | 370  2900  630  540  -  - | 11±0.06 | 6100  710  0  0  200  1300 | 14±0.1 | 570  2500  -  -  230  350 | 9±0.04 |
| β-1,3-glucan | C1  C2 | 103.5  74.4 | 176.2  176.3 | 420  400 | 4±0.03 | 620  420 | 3±0.04 | 480  340 | 4±0.02 | 730  620 | 7±0.02 | 2100  1700 | 13±0.05 | 1400  1100 | 12±0.1 |
| Galactose | C1  C2  C2  C3  C3  C4  C4  C5  C5  C6 | 92.7  72.2  72.1  70.6  70.7  73.5  73.8  72.1  72.2  61.7 | 164.3  164.4  142.2  142.3  143.8  143.8  145.4  145.7  133.6  133.4 | 4000  3800  -  -  -  -  390  3100  2700  2000 | 29±0.1 | 4800  3500  -  -  -  -  5300  300  300  3000 | 26±0.2 | 4100  3300  3800  4500  -  -  4200  3300  1500  2700 | 37±2 | 1900  1600  -  -  -  -  1500  850  1900  2400 | 17±0.04 | 2100  5200  -  -  -  -  4800  7100  4200  2000 | 28±0.1 | 5000  4100  3600  3900  1700  1100  -  -  1600  3300 | 29±1 |
| Chitosan | C1  C2 | 101.8  55.7 | 157.6  158.3 | -  - |  | -  - |  | -  - |  | -  - |  | 810  790 | 5±0 | 560  440 | 5±0 |
| GalN | C1  C2 | 54.9 | 146.6 | -  - |  | -  - |  | -  - |  | 1000  1100 | 9±0.01 | -  - | -  - | 230  170 | 2±0 |
|  |  | 91.7 | 146.1 |  |  |  |  |  |  |  |  |  |  |  |  |
| GalNAc | C1  C2 | 95.6  57.52 | 152.7  152.9 | -  - |  | -  - |  | -  - |  | -  - | 6±0 | -  - | -  - | 767  671 | 7±0.01 |

**Table S6. Water-edited intensities of the polysaccharides**. The intensity ratios are obtained by comparing the peak intensities between water-edited and control 2D spectra of *A. atacamensis* cell walls. The error bars represent propagated standard deviations from NMR signal-to-noise ratios, with an error margin typically below 10%.

| **Cross peaks** | **1d, 5.13M NaCl** | **4d, 5.13M NaCl** | **7d, 1.7M NaCl** | **10d, 0.5M MgCl_2_** | **10d, 1.5M MgCl_2_** | **Cross peaks** | **1d, 5.13M NaCl** | **4d, 5.13M NaCl** | **7d, 1.7M NaCl** | **10d, 0.5M MgCl_2_** | **10d, 1.5M MgCl_2_** |
| --- | --- | --- | --- | --- | --- | --- | --- | --- | --- | --- | --- |
| B1-1 | 0.25±0.06 | 0.13±0.04 | 0.38±0.06 | 0.80±0.03 | 0.75±0.07 | Ch1-1 | 0.25±0.05 | 0.13± 0.04 | 0.38±0.07 | 0.75±0.03 | 0.75±0.07 |
| B1-3 | 0.5±0.2 | 0.21±0.10 | 0.5±0.2 | - | 0.13±0.05 | Ch1-3 | - | - | - | 0.5±0.1 | 0.6±0.2 |
| B1-5 | 0.4±0.1 | - | 0.3±0.2 | - | 0.9±0.3 | Ch1-5 | - | 0.13±0.06 | - | - | - |
| B1-2 | 0.34±0.09 | 0.13±0.06 | 0.4±0.1 | - | 2.0±0.1 | Ch1-3 | 0.6±0.2 | - | 0.3±0.1 | - | - |
| B1-4 | - | - | 0.4±0.2 | 0.50±0.05 | 0.13±0.05 | Ch1-2 | - | - | - | 0.43±0.08 | 0.4±0.1 |
| B3-1 | 0.5±0.2 | - | 0.4±0.1 | 0.8±0.07 | - | Ch4-1 | - | - | - | 0.5±0.1 | 0.3±0.1 |
| B3-3 | 0.4±0.1 | 0.15±0.07 | 0.4±0.1 | 0.8±0.09 | 0.9±0.2 | Ch4-4 | - | 0.3±0.1 | - | 0.42±0.08 | 0.4±0.1 |
| B3-5 | 0.4±0.2 | - | 0.5±0.2 | - | - | Ch4-5 | - | - | - | 0.3±0.1 | 0.4±0.1 |
| B3-2 | 0.5±0.2 | 0.18±0.07 | 0.5±0.2 | 0.90±0.08 | 0.9±0.2 | Ch4-3 | - | - | - | 0.41±0.09 | 0.6±0.1 |
| B3-4 | 0.4±0.1 | 0.14±0.08 | 0.6±0.2 | 0.90±0.09 | 0.8±0.2 | Ch4-6 |  | - | - | 0.5±0.1 | 0.3±0.2 |
| B3-6 | 0.3±0.2 | - | 0.6±0.3 | - | 0.9±0.3 | Ch4-2 | 0.28±0.04 | - | - | 0.4±0.1 | 0.4±0.1 |
| B5-1 | - | 0.22±0.06 | 0.4±0.2 | - | 0.6±0.2 | Ch5-1 | - | - | 0.3±0.1 | - | 0.8±0.1 |
| B5-3 | 0.4±0.2 | - | 0.4±0.2 | - | - | Ch5-3 | - | - | - | - | 0.6±0.2 |
| B5-5 | 0.43±0.06 | 0.17±0.03 | 0.41±0.06 | 0.75±0.04 | 0.65±0.07 | Ch5-5 | - | - | 0.34±0.04 | - | 0.55±0.04 |
| B5-2 | 0.4±0.1 | 0.14±0.04 | - | 0.8±0.07 | 0.7±0.1 | Ch5-6 | - | - | 0.4±0.1 | - |  |
| B5-4 | 0.4±0.1 | 0.10±0.05 | 0.4±0.1 | - | 0.9±0.2 | Ch5-2 | - | - | - | - | 0.7±0.2 |
| B5-6 | 0.4±0.1 | 0.17±0.04 | 0.5±0.1 | 0.93±0.06 | - | Ch3-1 | - | - | - | 0.13±0.01 | 0.8±0.2 |
| B2-1 | 0.33±0.08 | 0.14±0.03 | 0.5±0.1 | 0.90±0.04 | - | Ch1-4 | 0.5±0.2 | - | - | 0.6±0.1 | 0.4±0.2 |
| B2-3 | 0.5±0.2 | 0.16±0.06 | 0.4±0.2 | - | 0.12±0.03 | Ch3-5 | 0.3±0.1 |  | 0.3±0.1 | - | - |
| B2-5 | 0.5±0.2 | 0.18±0.04 | 0.3±0.2 | 0.92±0.09 | - | Ch3-3 | 0.49±0.07 | 0.25±0.04 | 0.67±0.06 | 0.64±0.03 | 0.42±0.03 |
| B2-2 | 0.4±0.05 | 0.18±0.02 | 0.51±0.05 | 0.76±0.02 | 0.67±0.04 | Ch3-6 | - | 0.2±0.1 | - | 1.0±0.1 | - |
| B2-4 | 0.4±0.1 | 0.12±0.06 | - | - | - | Ch3-2 | - | - | 0.4±0.2 | 0.44±0.06 | 0.4±0.1 |
| B2-6 | 0.3±0.1 | - | 0.5±0.2 | 0.83±0.07 | 0.7±0.1 | Ch2-1 | - | - | - | 0.41±0.07 | 0.5±0.1 |
| B4-1 | 0.5±0.1 | 0.15±0.05 | 0.4±0.1 | 0.44±0.08 | - | Ch2-4 | - | - | - | 0.4±0.1 | 0.5±0.2 |
| B4-3 | 0.4±0.2 | 0.20±0.08 | 0.4±0.1 | 0.4±0.1 | - | Ch2-5 | - | - | - | 0.5±0.1 | 0.6±0.1 |
| B4-5 | - | 0.19±0.07 | 0.4±0.1 | 0.32±0.07 | - | Ch3-3 | - | - | - | 0.42±0.06 | 0.3±0.1 |
| B4-2 | 0.5±0.2 | 0.23±0.07 | 0.4±0.1 | 0.45±0.07 | - | Ch3-6 | - | - | - | 0.5±0.1 | 0.5±0.2 |
| B4-4 | 0.43±0.07 | 0.18±0.04 | 0.39±0.09 | 0.43±0.05 | 0.86±0.08 | Ch3-2 | 0.4±0.2 | 0.3±0.1 | 0.3±0.1 | 0.40±0.04 | 0.4±0.1 |
| B4-6 | - | 0.17±0.06 | 0.5±0.2 | 0.24±0.01 | - | **Average**  **n** | **0.40**  **7** | **0.22**  **6** | **0.38**  **9** | **0.48**  **20** | **0.51**  **23** |
| **Average**  **n** | **0.41**  **25** | **0.17**  **22** | **0.48**  **27** | **0.68**  **19** | **0.75**  **17** | Cs1-2  Cs1-3  Cs1-5 | **-**  **-**  **-** | **-**  **-**  **-** | **-**  **-**  **-** | 0.33±0.1  0.55±0.06  0.40±0.07 | 0.3±0.02  0.48±0.08  0.81±0.06 |
|  |  |  |  |  |  | **Average** |  |  |  | **0.43** | **0.53** |

**Table S7.^13^C-T_1_ relaxation times and ^1^H-T_1ρ_ relaxation times of polysaccharides.** T_1_ relaxation time constants were determined by fitting the data to a single-exponential equation function: $I\left( t \right)=e^{-t/T_{I}}$ . Similarly, T_1ρ_ relaxation times were obtained using the single exponential equation $I\left( t \right)=e^{-t/T_{I\rho}}$. Error bars are standard deviations of the fit parameters.

| Glucans Carbons sites/  Sample | ^13^C-T_1_ (s) | |  | Glucans Carbons /  Sample | ^1^H-T_1ρ_ (ms) | |
| --- | --- | --- | --- | --- | --- | --- |
|  | **MgCl_2_** | |  |  | **MgCl_2_** | |
|  | **10 d, 1.5 M** | **10 d, 0.5 M** |  |  | **10 d, 1.5 M** | **10 d, 0.5 M** |
| B1 | 2.6±0.3 | 2.4±0.2 |  | B1 | 10.0±0.8 | 9.5±0.8 |
| B3 | 1.2±0.1 | 1.4±0.1 |  | B3 | 11.2±0.6 | 6.9±0.6 |
| B5 | 1.4±0.2 | 1.4±0.1 |  | B5 | 11.4±1.0 | 6.4±0.6 |
| B2 | 2.0±0.2 | 1.8±0.2 |  | B2 | 5.6±0.4 | 7.7±0.8 |
| B4  B6 | 1.1±0.1  1.4±0.2 | 1.2±0.1  1.4±0.2 |  | B4  B6 | 7.1±0.5  6.3±0.5 | 6.1±0.6  5.9±0.6 |
| Average | **1.6** | **1.6** |  | **Average** | **8.6** | **7.1** |
| Ch1 | 2.6±0.3 | 2.4±0.2 |  | Ch1 | 10.1±0.7 | 9.5±0.8 |
| Ch4 | 4.2±0.3 | 3.9±0.3 |  | Ch4 | 14.1±1.0 | 12.8±1.1 |
| Ch3 | 2.7±0.3 | 2.4±0.3 |  | Ch3 | 9.8±0.7 | 8.7±0.8 |
| Ch2 | 2.6±0.3 | 2.9±0.3 |  | Ch2 | 9.1±0.8 | 9.7±0.8 |
| Average | **3.0** | **2.9** |  | **Average** | **10.8** | **10.2** |
| NaCl Conditions | | | | | | |
| Sample | **7d, 1.7 M** | **1d, 5.13 M** |  | **Sample** | **7d, 1.7 M** | **1d, 5.13 M** |
| B1 | 2.1±0.1 | 1.9±0.1 |  | B1 | 12.1±0.5 | 10.2±0.6 |
| B3 | 1.3±0.1 | 1.3±0.1 |  | B3 | 9.4±0.6 | 12.0±0.9 |
| B5 | 1.5±0.1 | 1.3±0.1 |  | B5 | 9.7±0.6 | 11.0±0.7 |
| B2 | 1.9±0.1 | 1.6±0.1 |  | B2 | 10.5±0.6 | 9.9±0.7 |
| B4 | 1.4±0.1 | 1.3±0.1 |  | B4 | 9.9±0.4 | 8.3±0.6 |
| B6 | - | 1.3±0.1 |  | B6 | - | 8.4±0.6 |
| Average | **1.6** | **1.5** |  | **Average** | **10.3** | **10.0** |
| Ch1 | 2.1±0.1 | 1.9±0.1 |  | Ch1 | 12.1±0.5 | 10.2±0.6 |
| Ch4 | 3.7±0.2 | 4.0±0.3 |  | Ch4 | 13.7±0.5 | - |
| Ch5 | 2.4±0.2 | 2.1±0.2 |  | Ch5 | - | - |
| Ch3 | 2.6±0.2 | 2.0±0.2 |  | Ch3 | 10.6±0.8 | 9.6±0.6 |
| Ch2 | 3.3±0.2 | 2.3±0.2 |  | Ch2 | 10.8±0.6 | 9.2±0.7 |
| Average | **2.8** | **2.5** |  | **Average** | **11.8** | **9.7** |
| Sample | **4d, 5.13 M** | |  | **Sample** | **4d, 5.13 M** | |
| B1 | 1.6±0.2  1.2±0.1  1.1±0.1  1.4±0.2  1.1±0.1  1.1±0.1 | |  | B1 | 9.8±0.5  9.3±0.6  8.4±0.7  8.5±0.5  7.4±0.4  7.6±0.5 | |
| B3 |  |  |  | B3 |  |  |
| B5 |  |  |  | B5 |  |  |
| B2 |  |  |  | B2 |  |  |
| B4 |  |  |  | B4 |  |  |
| B6 |  |  |  | B6 |  |  |
| Average | **1.25** | |  | **Average** | **8.5** | |
| Ch1 | 1.6±0.2  2.5±0.4  -  1.7±0.2  2.0±0.3 | |  | Ch1 | 9.8±0.5  11.6±0.7  -  8.0±0.4  7.9±0.5 | |
| Ch4 |  |  |  | Ch4 |  |  |
| Ch5 |  |  |  | Ch5 |  |  |
| Ch3 |  |  |  | Ch3 |  |  |
| Ch2 |  |  |  | Ch2 |  |  |
| Average | **2.0** | |  | **Average** | **9.3** | |

14.	Dickwella Widanage, M. C.; Gautam, I.; Sarkar, D.; Mentink-Vigier, F.; Vermaas, J. V.; Ding, S.-Y.; Lipton, A. S.; Fontaine, T.; Latgé, J.-P.; Wang, P., Adaptative survival of Aspergillus fumigatus to echinocandins arises from cell wall remodeling beyond β− 1, 3-glucan synthesis inhibition. *Nat. Commun.* **2024,** *15*, 6382.

15.	Fontaine, T.; Delangle, A.; Simenel, C.; Coddeville, B.; van Vliet, S. J.; Van Kooyk, Y.; Bozza, S.; Moretti, S.; Schwarz, F.; Trichot, C., Galactosaminogalactan, a new immunosuppressive polysaccharide of Aspergillus fumigatus. *PLoS Pathog.* **2011,** *7*, e1002372.

16.	Kirui, A.; Zhao, W.; Deligey, F.; Yang, H.; Kang, X.; Mentink-Vigier, F.; Wang, T., Carbohydrate-aromatic interface and molecular architecture of lignocellulose. *Nature communications* **2022,** *13* (1), 538.

17.	Latgé, J. P.; Kobayashi, H.; Debeaupuis, J. P.; Diaquin, M.; Sarfati, J.; Wieruszeski, J. M.; Parra, E.; Bouchara, J. P.; Fournet, B., Chemical and Immunological Characterization of the Extracellular Galactomannan of Asperillus fumigatus. *Infect. Immun.* **1994,** *62*, 5424-5433.

18.	Chakraborty, A.; Fernando, L. D.; Fang, W.; Dickwella Widanage, M. C.; Wei, P.; Jin, C.; Fontaine, T.; Latgé, J. P.; Wang, T., A molecular vision of fungal cell wall organization by functional genomics and solid-state NMR. *Nat. Commun.* **2021,** *12*, 6346.

19.	Poulhazan, A.; Dickwella Widanage, M. C.; Muszyński, A.; Arnold, A. A.; Warschawski, D. E.; Azadi, P.; Marcotte, I.; Wang, T., Identification and Quantification of Glycans in Whole Cells: Architecture of Microalgal Polysaccharides Described by Solid-State Nuclear Magnetic Resonance. *J. Am. Chem. Soc.* **2021**.

20.	Fernando, L. D.; Pérez-Llano, Y.; Dickwella Widanage, M. C.; Jacob, A.; Martínez-Ávila, L.; Lipton, A. S.; Gunde-Cimerman, N.; Latgé, J.-P.; Batista-García, R. A.; Wang, T., Structural adaptation of fungal cell wall in hypersaline environment. *Nat. Commun.* **2023,** *14* (1), 7082.
